# DEAD box RNA helicase 5 is a new pro-viral host factor for Sindbis virus infection

**DOI:** 10.1101/2023.09.21.558232

**Authors:** Mélanie Messmer, Louison Pierson, Charline Pasquier, Nikola Djordjevic, Johana Chicher, Philippe Hammann, Sébastien Pfeffer, Erika Girardi

## Abstract

**Background:** RNA helicases are emerging as key factors regulating host-virus interactions. The DEAD-box ATP-dependent RNA helicase DDX5, which plays an important role in many aspects of cellular RNA biology, was also found to either promote or inhibit viral replication upon infection with several RNA viruses. Here, our aim is to examine the impact of DDX5 on Sindbis virus (SINV) infection.

**Methods:** We analysed the interaction between DDX5 and the viral RNA using imaging and RNA-immunoprecipitation approaches. The interactome of DDX5 in mock- and SINV-infected cells was determined by mass spectrometry. We validated the interaction with DDX17 and the viral capsid by co-immunoprecipitation in the presence or absence of an RNase treatment. We determined the subcellular localization of DDX5, its cofactor DDX17 and the viral capsid protein by co-immunofluorescence. Finally, we investigated the impact of DDX5 depletion and overexpression on SINV infection at the viral protein, RNA and infectious particle accumulation level. The contribution of DDX17 was also tested by knockdown experiments.

**Results:** In this study we demonstrate that DDX5 interacts with the SINV RNA during infection. Furthermore, the proteomic analysis of the DDX5 interactome in mock and SINV-infected HCT116 cells identified new cellular and viral partners and confirmed the interaction between DDX5 and DDX17. Both DDX5 and DDX17 re-localize from the nucleus to the cytoplasm upon SINV infection and interact with the viral capsid protein. We also show that DDX5 depletion negatively impacts the viral replication cycle, while its overexpression has a pro-viral effect. Finally, we observed that DDX17 depletion reduces SINV infection, an effect which is even more pronounced in a DDX5-depleted background, suggesting a synergistic pro-viral effect of the DDX5 and DDX17 proteins on SINV.

**Conclusions:** These results not only shed light on DDX5 as a novel and important host factor to the SINV life cycle, but also expand our understanding of the roles played by DDX5 and DDX17 as regulators of viral infections.

## 1. Background

DEAD box RNA helicases play crucial molecular functions in virtually all aspects of RNA metabolism, including transcription, pre-mRNA splicing, RNA transport, rRNA biogenesis and RNA decay (1). These enzymes can structurally remodel short RNA duplex molecules and rearrange RNA-protein interactions in an ATP-dependent fashion. The DEAD-box RNA helicase family is characterized by the presence of specific highly conserved motifs, including the Asp–Glu–Ala–Asp (D-E-A-D) box within the catalytic core (2) from which this family gets its name. Recently, DEAD box RNA helicases have emerged as pivotal host factors upon viral infection that act either by modulating antiviral signalling pathways or by directly recognizing viral RNA and acting at different steps of the viral life cycle (3–5).

The multifunctional DEAD-box RNA helicase DDX5 (also known as p68) is an essential splicing factor (6,7) involved in many other aspects of cellular RNA metabolism (8). Within the DDX gene family members, DDX5 exhibits the highest degree of homology with DDX17 (also known as p72) (9). These two proteins can form homodimers but are preferentially associated in hetero-dimers (10). DDX5 and DDX17 exhibit partially overlapping functions, including involvement in transcriptional regulation (11), alternative splicing modulation (12) and ribosomal biogenesis (13).

A growing body of evidence suggests that DDX5 and DDX17 play crucial roles during viral infection (14,15). Despite their known function as a heterodimer, these two proteins exhibit distinct functions in response to specific viral infections. For instance, depletion of DDX17 but not of DDX5 increases infection of Rift Valley Fever virus (RVFV), a segmented, negative-sense RNA virus. In particular, DDX17 binding to a stem-loop structure in the viral RNA is sufficient to trigger the antiviral effect (16).

Moreover, the same protein can have either a positive or a negative effect on different viruses. While DDX5 is antiviral in response to DNA viruses such as hepatitis B virus (HBV) (17) or myxoma virus (MYXV) (18), it acts as a positive cofactor of the Human Immunodeficiency Virus 1(HIV-1) proteins Rev and Tat facilitating viral replication (19,20). Interestingly, several positive, single-stranded RNA viruses with a cytoplasmic replication cycle have also evolved to hijack DDX5 and favour their own replication. DDX5 was also shown to be pro-viral for hepatitis C virus (HCV) (21,22), severe acute respiratory syndrome coronavirus (SARS-CoV) (23), japanese encephalitis virus (JEV) (24) and porcine reproductive and respiratory syndrome virus (PRRSV) (25). In most cases, DDX5 promotes viral replication through direct interaction with specific viral proteins and/or viral RNA.

In a previous study we identified DDX5 as one of the factors associated with the Sindbis virus (SINV) replication complex by dsRNA-IP coupled to mass spectrometry (DRIMS) and observed that DDX5 knockdown (KD) significantly reduces GFP expression in HCT116 cells infected with a GFP-expressing SINV (26). SINV is a small, enveloped, arthropod-borne virus, which belongs to the *Alphavirus* genus from the *Togaviridae* family. Alphaviruses are widely distributed viruses (27) that pose a re-emerging threat to human health due to their potential to cause severe arthritogenic and neurological diseases and for which approved vaccines are not yet available. Most of our understanding on the alphavirus life cycle derives from research on the prototypical model SINV. The virus possesses a positive, single-stranded genomic RNA of approximately 11 kb, capped and polyadenylated, organized into two distinct open reading frames (ORFs). The first ORF encodes the non-structural proteins (nsP1, nsP2, nsP3 and nsP4) necessary for viral RNA replication. An antigenomic RNA accumulates in the cytoplasm and serves as a template for the synthesis of both genomic and sub-genomic RNA. The second ORF is translated from the sub-genomic RNA and gives rise to the structural proteins (Capsid, E3, E2, 6K and E1) needed for viral particle assembly (28).

In this study we investigated the impact of DDX5 on SINV. Initially, we observed the co-localization and interaction between DDX5 and the viral RNA in the cytoplasm of infected cells by imaging and RNA IP approaches. We then conducted a proteomic analysis to determine the interactome of DDX5 under both uninfected and SINV-infected conditions. DDX17, a previously well-known DDX5 partner, emerged among the most significant hits. Interestingly, we demonstrated that DDX5 and DDX17 re-localize to the cytoplasm upon SINV infection and identified a novel RNA-independent interaction between DDX5 and viral capsid proteins.

Functionally, our study demonstrated that reducing DDX5 levels by either knock-down or knock-out approaches diminishes SINV replication in human HCT116 cells. Moreover, stable overexpression of DDX5 in *DDX5* knock-out cells restored the capsid protein expression and the viral particle production. Finally, we showed that knockdown of DDX17 has a negative impact on SINV viral cycle with a more pronounced decrease in cells lacking both DDX17 and DDX5, underscoring the importance of both DDX5 and DDX17 in promoting SINV infection.

## 2. Methods

### 2.1. Cell lines and viruses

HCT116, BHK21 and Vero E6 cells were maintained at 37°C in a humidified atmosphere containing 5% CO_2_ and cultured in Dulbecco’s Modified Eagle medium (DMEM; Invitrogen) supplemented with 10% FBS. Viral stocks were prepared from infectious clones (kindly provided by Dr Carla Saleh, Institut Pasteur, Paris, France) carrying either the wild-type or a GFP containing version of the SINV genomic sequence as previously described in (29). Briefly, the plasmids were linearized by XhoI digestion and used as a substrate for *in vitro* transcription using the mMESSAGE mMACHINE capped RNA transcription kit (Ambion, Thermo Fisher Scientific Inc.). In vitro synthesized viral RNA was transfected in BHK21 cells, viral infectious particles were collected 48 hpt and used to infect BHK21 for the full viral production. Titers were measured by standard plaque assay in Vero E6 cells. Cells were infected at a MOI of 1, 10^-1^ or 10^-2^ as indicated in the figure legends.

### 2.2. Plaque assay

For plaque assay, 1 million Vero E6 cells were seeded in a 96 well plate and infected for 1 hour with viral supernatants prepared in 10-fold dilution cascade. After removal of the viral inoculum, 2.5% carboxymethyl cellulose diluted in DMEM-FBS10% was added and cells were incubated at 37°C in a humidified atmosphere of 5% CO2. Plaques were counted manually under the microscope after 72 h.

### 2.3. DDX5 CRISPR/Cas9 knockout

Guide RNA (gRNA) sequences targeting the human DDX5 gene were designed using the CRISPOR tool (http://crispor.tefor.net) and cloned into the pX459-V2 vector (Addgene Plasmid #62988 was a gift from Feng Zhang (30)). Briefly 0.25 µM of the annealed oligos and 100 ng of digested vector were ligated and transformed into DH5 alpha cells.

guideRNA#2 DDX5 sense: 5’ CACCGATAATAGGGTGTTCATAGGT 3’
guideRNA#2 DDX5 antisense: 5’ AAACACCTATGAACACCCTATTATC 3’
guideRNA#1 DDX5 sense: 5’ CACCGCCCTACTTCCTCCAAATCG 3’
guideRNA#1 DDX5 antisense: 5’ AAACCGATTTGGAGGAAGTAGGGC 3’

HCT116 cells were transfected with 2 plasmids containing the sequence of gRNA#1 and #2, respectively. Twenty-four hours post-transfection, cells were treated with 1 µg/mL puromycin for 48h. Surviving cells were diluted to obtain 0.5 cells/ well in 96 well plates. Two weeks later, cellular genomic DNA was extracted from individual colonies. Cells were lysed in genomic DNA extraction buffer (50 mM Tris-HCl [pH 8.0]; 100 mM EDTA [pH 8.0], 100 mM NaCl, 1% SDS) containing 0.1 mg of proteinase K and incubated overnight at 55°C. Then, 50 ng of genomic DNA were amplified with the GoTaq (Promega) using specific primers (IDT) to detect the deletion (DDX5 FW: 5’-ATAAATCCCCGGCTTCCGAC-3’; DDX5 RV: 5’-AGAGGGGGTAGGTGGAAACAA-3’). Wild type genomic DNA was used as control template. PCR reactions were loaded on a 1 % agarose gel and the obtained amplicons were gel purified and sequenced by Sanger sequencing.

### 2.4. SiRNA-based knockdown

For the knockdown experiments, 0.6 millions of cells HCT116 cells were reverse-transfected with 20nM of ON-TARGETplus Human siRNAs (Horizon) against DDX5 (L-003774-0-0005), DDX17 (L-013450-01-0005) or non-targeting Control Pool (D-001810-10-05) using the Lipofectamine 2000 transfection reagent (Thermo Fisher Scientific, #11668019) according to the manufacturer’s instructions in 6 well plates. After 24 hours, cells were transfected again with 20nM of the same mix of siRNAs and incubated overnight. Cells were then infected with SINV-GFP at a MOI of 0.1 for 24 hours. Supernatant were collected for plaque assay experiment and proteins and RNA were collected for western blot and RT-qPCR analyses, respectively.

### 2.5. Lentivirus production and generation of stable cell lines

Lentiviral particles were produced by transfecting seven millions of HEK293T cells with 12 µg of either pLV-DDX5-V5 or pLV-BFP viral vector (Vector Builder), 2.4 µg and 9,6µg of the packaging plasmids pCMV-VSV-G (Addgene #8454) and psPAX2 (Addgene #12260) respectively, using Lipofectamine 2000 reagent (Invitrogen, Fisher Scientific) according to the manufacturer’s protocol. The viral supernatants were collected after 48 hours and filtered using a 0.45µm PES filter.

*DDX5* KO HCT116 cells were transduced in 6-well plates using the lentiviral supernatant diluted in fresh medium supplemented with 4μg/mL of polybrene (Merck, Sigma-Aldrich). The medium was changed after 8 h and the selection with the appropriate antibiotic was initiated after 48 h. Polyclonal V5-DDX5-*DDX5* KO HCT116 (V5-DDX5) and BFP-*DDX5* KO HCT116 (BFP) cells were maintained under selection for a minimum of 10 days before analysis. In detail, two-hundred thousand of V5-DDX5 or BFP-DDX5 stable cells were infected with SINV-GFP for 24 hours (MOI of 0.1). before collecting the supernatant for plaque assay experiment, as well as proteins and RNA for western blot and RT-qPCR analyses, respectively.

### 2.6. Live-cell imaging

Two-hundred thousand WT, *DDX5* KO, BFP or V5-DDX5-HCT116 cells were seeded in 12 well plate and infected with SINV-GFP at the indicated MOI. Uninfected cells were used as control. GFP fluorescence and phase contrast were observed using a CellcyteX live-cell imaging system (Discover Echo). Four to six images per well (10X objective) were acquired every 3 or 6 hours for 72hrs and were analysed with the Cellcyte Studio software to determine cell confluency and GFP relative intensity.

### 2.7. Cloning

Amplicons containing the coding sequence for the individual viral proteins were generated by PCR from the plasmid carrying the SINV genomic sequence (kindly provided by Dr. Carla Saleh, Institut Pasteur, Paris, France) and cloned into the pDONOR221 vector (Invitrogen) by recombination using the Gateway BP clonase (Invitrogen) according to the manufacturer instructions. The pDEST-FLAG-HA-nsP1, pDEST-FLAG-HA-nsP2, pDEST-FLAG-HA-nsP3, pDEST-FLAG-HA-nsP4 and pDEST-FLAG-HA-capsid vectors were obtained by recombination using Gateway LR clonase (Invitrogen) in the pDEST-FLAG-HA vector according to the manufacturer instructions, as in (31).Primers used for the PCR are listed below:

pDONOR NSP1 FW
5’ GGGGACAAGTTTGTACAAAAAAGCAGGCTTCGAGAAGCCAGTAGTAAAC-3’
pDONOR NSP1 REV
5’-GGGGACCACTTTGTACAAGAAAGCTGGGTCTTATTTGCTCCGATGTCCG-3’
pDONOR NSP2 FW
5’GGGGACAAGTTTGTACAAAAAAGCAGGCTTCGCATTAGTTGAAACCCCG-3’
pDONOR NSP2 REV
5’GGGGACCACTTTGTACAAGAAAGCTGGGTCTTATTGGCTCCAACTCCATCTC-3’
pDONOR NSP3 FW
5’-GGGGACAAGTTTGTACAAAAAAGCAGGCTTCGCGCCGTCATACCGCACC-3’
pDONOR NSP3 REV
5’-GGGGACCACTTTGTACAAGAAAGCTGGGTCTTATTGTATTCAGTCCTCC-3’
pDONOR NSP4 FW
5’-GGGGACAAGTTTGTACAAAAAAGCAGGCTTCCTAACCGGGGTAGGTGGGTAC 3’
pDONOR NSP4 REV
5’GGGGACCACTTTGTACAAGAAAGCTGGGTCTTACTATTTAGGACCACCGTAGAG -3’
pDONOR capsid FW
5’-GGGGACAAGTTTGTACAAAAAAGCAGGCTTCAATAGAGGATTCTTTAAC-3’
pDONOR capsid REV
5’-GGGGACCACTTTGTACAAGAAAGCTGGGTCTTATTCCACTCTTCTGTCC-3’

### 2.8. Flag-HA SINV protein co-IP

For the experiment, 0.6 millions of HCT 116 cells were seeded in 6 well plates. The following day 3µg of the pDEST-HA-FLAG described above were transfected using the Lipofectamine 2000 transfection reagent as per manufacturer’s recommendations. 48 hours later, the cells were washed once with 1xPBS and lysed in 600 µL of Miltenyi lysate buffer (50 mM Tris-HCl pH 7.5, 140 mM NaCl, 1.5 mM MgCl_2_, 0.1% NP-40), supplemented with Complete-EDTA-free Protease Inhibitor Cocktail (Merck). Cells were lysed for 30 min incubation on ice and debris were removed by 10 min centrifugation at 10,000g at 4°C. An aliquot of the cleared lysates (50 μL) was kept aside as protein Input. Samples were divided into equal parts (250 μL each) and incubated with 50 μL of magnetic microparticles coated with monoclonal HA or MYC antibodies (MACS purification system, Miltenyi Biotec) at 4°C for 1 hour under rotation (10 rpm). Samples were passed through μ Columns (MACS purification system, Miltenyi Biotec). The μ Columns were then washed 4 times with 200 μL of WASH buffer1 and 1 time with 200 μL of WASH buffer 2. To elute the immunoprecipitated proteins, 70µL of 95°C pre-warmed 2x Western blot loading buffer (10% glycerol, 4% SDS, 62.5 mM Tris-HCl pH 6.8, 5% (v/v) 2-β-mercaptoethanol, Bromophenol Blue) was passed through the μ Columns. Proteins were analysed by western blotting.

### 2.9. DDX5 and DDX17 co-immunoprecipitation and RNA immunoprecipitation

DDX5 and DDX17 immunoprecipitations (IP) were performed as in (26) with some modifications. Briefly, 4 millions of mock or SINV-GFP infected HCT116 cells (MOI 0.1, 24 hpi) were lysed using RIP immunoprecipitation buffer (50mM tris-HCL [pH 7.5], 150mM NaCl, 5 mM EDTA, 0.05% SDS, 1% triton, 1 tablet of commercial protease inhibitor cocktail (Merck)). RNase treatment was performed on the lysates by adding 1µL of RNase A at 10 mg/mL (Thermo Fisher) and incubating for 25 min at 37°C. DNase treatment was performed on the lysates by adding 1µL of RNase-free DNase I at 1U/µL (Thermo Fisher Scientific), 10mM MgCl2, 5mM Cacl2 and 1µL of Ribolock (Thermo Fisher Scientific) to the 1 mL lysate and incubating for 20 min at 37°C. Lysates were centrifuged for 10 min at 4°C. Supernatants were collected and pre-cleared for 1h at room temperature with Dynabeads protein G magnetics beads (Invitrogen) blocked with yeast tRNA (Invitrogen). The efficiency of the RNase treatment was evaluated by RNA analysis of the input samples on a 1% agarose gel.

Lysates were incubated over night at 4°C with 40 µL of Dynabeads protein G beads conjugated with 2 µg rabbit anti-DDX5 antibody (ab21696; Abcam), or rabbit anti-DDX17 antibody (19910-1-AP, Proteintech) or 2µg rabbit IgG antibody (2729; Cell Signaling Technology).

Beads were washed 3 times with RIP buffer containing 150mM NaCl, twice with the same buffer supplemented with 50mM of NaCl, and 3 times 3 times with RIP buffer containing 150mM NaCl. 30% of the immunoprecipitated protein and 70% of the associated RNA were eluted with Laemmli 1X at 95°C for 10 min or with 1 mL TriReagent (Thermo Fisher Scientific), respectively. Proteins were analysed by western blot or by mass spectrometry, while RNA was extracted and analysed by RT-qPCR.

### 2.10. Western Blot

Cells were collected in 300 to 500 µL of lysis buffer (50mM tris-HCL [pH 7.5], 150mM NaCl, 5 mM EDTA, 0.05% SDS, 1% triton, 1 tablet of commercial protease inhibitor cocktail (ROCHE)) and incubated for 30 min on ice. Cell lysates were then collected and protein concentration was determined using the Bradford method (Bio-Rad). For total protein analysis, 20µg of protein samples were heated in 1X Laemmli buffer at 95°C for 5 min and loaded on pre-cast 4-20% SDS-polyacrylamide electrophoresis gel (Bio-Rad). For the RIP experiment, 0.5 % of input and 12,5% of immunoprecipitated protein were used. After migration, the proteins were transferred onto 0.45 μm nitrocellulose membranes (GE healthcare). The membranes were blocked with 5% non-fat milk diluted in PBS 1X (Euromedex) complemented with 0.2% Tween-20 (PBS-T) for 1 h at room temperature and then were incubated at 4°C overnight or 1 h at room temperature with the following specific primary antibodies: anti-DDX5 (mouse-horseradish peroxidase(HRP), sc-365164; Santa Cruz Biotechnology) or anti-DDX5 (rabbit, ab21696; Abcam), anti-DDX17 (mouse, sc-271112; Santa Cruz Biotechnology), anti-tubulin HRP (ab21058; Abcam), anti-GAPDH-HRP (mouse, ab9482; Abcam), anti-capsid (rabbit, kind gift of Diane Griffin), anti-GFP (mouse, 11814460001, Roche). Membranes were washed 3 times with PBS-T for 5 min and incubated with mouse-HRP (A4416; Merck) and rabbit-HRP (GENA9640V, Merck) secondary antibodies for 1 h at room temperature. Proteins were detected using the Chemiluminescence substrate (Supersignal West Pico; Pierce) and analysed with the Fusion FX imaging system (Vilber).

### 2.11. RNA extraction and RT-qPCR

Total RNA and RNA enriched upon specific protein immunoprecipitation (RIP) were extracted using TriReagent (Invitrogen, Thermo Fisher Scientific) according to the manufacturer’s instructions. For total RNA analysis, 1 μg of RNA was reverse transcribed with the SuperScript VILO Master Mix (Invitrogen, Thermo Fisher Scientific) according to the manufacturer’s instructions. For the RIP experiment, 1 µL of samples before and after IP was used for retro-transcription using the same protocol. RT-qPCR was performed on cDNA using the Maxima SYBR Green qPCR Master Mix K0253; Thermo Fisher Scientific) on the CFX96 touch Real-Time PCR machine (Bio-Rad) using the following primers at an annealing temperature of 60°C:

SINV genome FW:
5^′^-CCACTACGCAAGCAGAGACG-3^′^;
SINV genome RV:
5^′^-AGTGCCCAGGGCCTGTGTCCG-3^′^;
GAPDH FW:
5^′^-CTTTGGTATCGTGGAAGGACT-3^′^;
GAPDH RV:
5^′^-CCAGTGAGCTTCCCGTTCAG-3’
SINV sub-genome-genome FW
5’CCACAGATACCGTATAAGGCA 3’
SINV sub-genome-genome RV
5’ TGCAGGTAATGTACTCTTGG 3’

### 2.12. Immunostaining

Mock or SINV WT-infected cells were plated on 8 wells LabTek slide (Merck Millipore), were fixed with 4% formaldehyde (Merck) diluted in PBS 1X for 10 min at room temperature and then washed 3 times with PBS 1X. Cells were blocked in blocking buffer (0.1% Triton X-100; PBS 1 X; 5% normal goat serum) for 1 h. The following primary antibodies were diluted 1:400 in blocking buffer and incubated over night at 4°C: mouse anti-dsRNA J2 (RNT-SCI-10010200; Jena bioscience), mouse anti-DDX5 (67025 Proteintech) or rabbit anti-DDX5 (ab21696; Abcam), anti-DDX17 (mouse, sc-27112; Santa Cruz Biotechnology), anti-capsid (rabbit, kind gift of Diane Griffin). Cells were washed with PBS 1X-Triton 0.1%. and incubated for 1 h at room temperature with secondary antibodies (goat anti-mouse Alexa 594 (A11032, Invitrogen) or goat anti-rabbit Alexa 488 (A11008, Invitrogen) secondary antibodies) diluted at 1:1000 in PBS 1X-Tween 0.1%. DAPI staining was performed for 5 min in PBS 1X to reveal the nuclei (D1306, Invitrogen, Thermo Fisher Scientific). Slides were mounted on coverslips with Fluoromount-G mounting media (Invitrogen, Thermo Fisher Scientific) and observed by confocal microscopy (LSM780, Zeiss) with a 40X or 63X objective. Images were analysed using Image J software and fluorescence intensity profiles were obtained.

### 2.13. Sequential immunostaining and FISH

Mock or SINV-WT infected (MOI 1, 24hpi) HCT116 cells were grown on 18 mm round cover glass in 12-well cell culture plates.

Cells were fixed with 3.7 % formaldehyde diluted in PBS 1X for 10 min at room temperature. Cells were then permeabilized in 1mL of 0.1% Triton X-100 in 1X PBS for 5 min at room temperature and incubated with anti-DDX5 primary antibody (rabbit, ab21696; Abcam) diluted to 1:400 in PBS 1X for 1 hour. Goat anti-rabbit Alexa 488 (A11008, Invitrogen) secondary antibody diluted to 1:1000 was added on the cells for 1 hour at room temperature. Cells were fixed again with 3.7 % formaldehyde (Biosearch technologies) diluted in PBS 1X for10 min at room temperature and incubated over night at room temperature with the SINV genome specific LGC Biosearch Technologies’ Stellaris® RNA FISH Probe diluted in RNA FISH hybridization buffer (Stellaris, Biosearch technologies). DAPI staining was performed for 30 min to reveal the nuclei (D1306, Invitrogen, Thermo Fisher Scientific). Slides were mounted on coverslips with Fluoromount-G mounting anti-fading media (Invitrogen, Thermo Fisher Scientific) and observed by confocal microscopy (LSM780, Zeiss). Images were analysed using Image J software.

### 2.14. Mass spectrometry analyses

For LC-MS/MS analyses, proteins were prepared as described in a previous study (26). Proteins eluted from the beads were washed with 2 sequential overnight precipitations with glacial 0.1 M ammonium acetate in 100% methanol (5 volumes) followed by 3 washes with glacial 0.1 M ammonium acetate in 80% methanol. Proteins were then solubilized in 50 mM ammonium bicarbonate for a reduction-alkylation step (dithiothreitol 5 mM – iodoacetamide 10 mM) and an overnight digestion with 300ng of sequencing-grade porcine trypsin (Promega, Fitchburg, MA, USA). Digested peptides were resuspended in 0.1% formic acid (solvent A) and injected on an Easy-nanoLC-1000 system coupled to a Q-Exactive Plus mass spectrometer (Thermo Fisher Scientific, Germany). One fourth of each sample was loaded on a C-18 precolumn (75 μm ID × 20 mm nanoViper, 3µm Acclaim PepMap; Thermo-Fisher Scientific) and separated on an analytical C18 analytical column (75 μm ID × 25 cm nanoViper, 3µm Acclaim PepMap) with a 160 minutes gradient of solvent B (0.1% of formic acid in acetonitrile).

Qexactive data were searched with Mascot algorithm (version 2.6.2, Matrix Science) against the Swissprot database with *H.sapiens* taxonomy (release 2020_05) as well as the sequences of GFP and of SINV virus (20 394 sequences) with a decoy strategy. The resulting .dat Mascot files were then imported into Proline v2.0 software (32) to align the identified proteins. Proteins were then validated on Mascot pretty rank equal to 1, 1% FDR on both peptide spectrum matches (PSM) and protein sets (based on Mascot score).

For statistical analyses of the mass spectrometry data, spectral counts (SpC) of the identified proteins were stored in a local MongoDB database and subsequently analysed through a Shiny Application built upon the R packages msmsEDA (Gregori J, Sanchez A, Villanueva J (2014). msmsEDA: Exploratory Data Analysis of LC-MS/MS data by spectral counts. R/Bioconductor package version 1.22.0) and msmsTests (Gregori J, Sanchez A, Villanueva J (2013). msmsTests: LC-MS/MS Differential Expression Tests. R/Bioconductor package version 1.22.0). Differential expression tests were performed using a negative binomial regression model. The P-values were adjusted with FDR control by the Benjamini–Hochberg method and the following criteria were used to define differentially expressed proteins: an adjusted P-value <0.05, a minimum of 5 SpC in the most abundant condition, and a minimum absolute Log2 fold change of 1.

### 2.15. Statistical analysis

Statistical analyses were performed using GraphPad Prism 9 software. Generally, a paired Student’s t test was performed for comparisons between two samples for plaque assays and a one-sample t test was performed to determine the significance of the fold change on qPCR results in which the control sample was set to 1. The number of replicates per experiment and the specific statistical tests used are stated in the figure legends.

## 3. Results

### 3.1. DDX5 interacts with SINV viral RNA in the cytoplasm of SINV -infected cells

It was previously reported that DDX5 is primarily localized in the nucleus (8), while the SINV replication cycle is strictly cytoplasmic (28). To determine whether DDX5 can affect SINV infection *via* a direct interaction with the viral components, we first verified whether DDX5 co-localizes with SINV RNA in infected cells. Despite the predominantly nuclear localization in uninfected cells, confocal microscopy analyses showed that DDX5 re-localizes to the cytoplasm in the presence of the virus and co-localizes with the viral genomic RNA at 24 hours post infection (**Figure 1A**), suggesting that DDX5 is recruited to the viral RNA. We subsequently performed RNA immunoprecipitation (RIP) on the endogenous DDX5 in mock and SINV GFP-infected HCT116 cells. Efficient DDX5 immunoprecipitation was assessed at the protein level by western blot (**Figure 1B**) and the associated RNA was analysed by RT-qPCR. We found that the viral genomic RNA is specifically enriched in the DDX5-RIP compared to the control (IgG-RIP) condition (**Figure 1C**). We also performed dot blot analyses using a J2 antibody, which detects dsRNA longer than 40 bp (33,34), and observed an enrichment of dsRNAs in the DDX5-IP compared to the IgG-IP in the infected samples (**Figure 1D**). Of particular note, we also observed enrichment of dsRNA in the mock condition, albeit to a lesser degree than upon SINV infection. This suggests that DDX5 binds to endogenous dsRNA molecules even in the absence of viral RNA. Whilst dsRNA can be found associated with DDX5 by dot blot analyses, confocal co-immunostaining analyses using the J2 antibody as a proxy for dsRNA do not demonstrate a clear co-localization of DDX5 and dsRNA upon SINV-infection (**Figure 1E**), but rather indicate a proximity between the protein and dsRNA.

**Figure 1:**
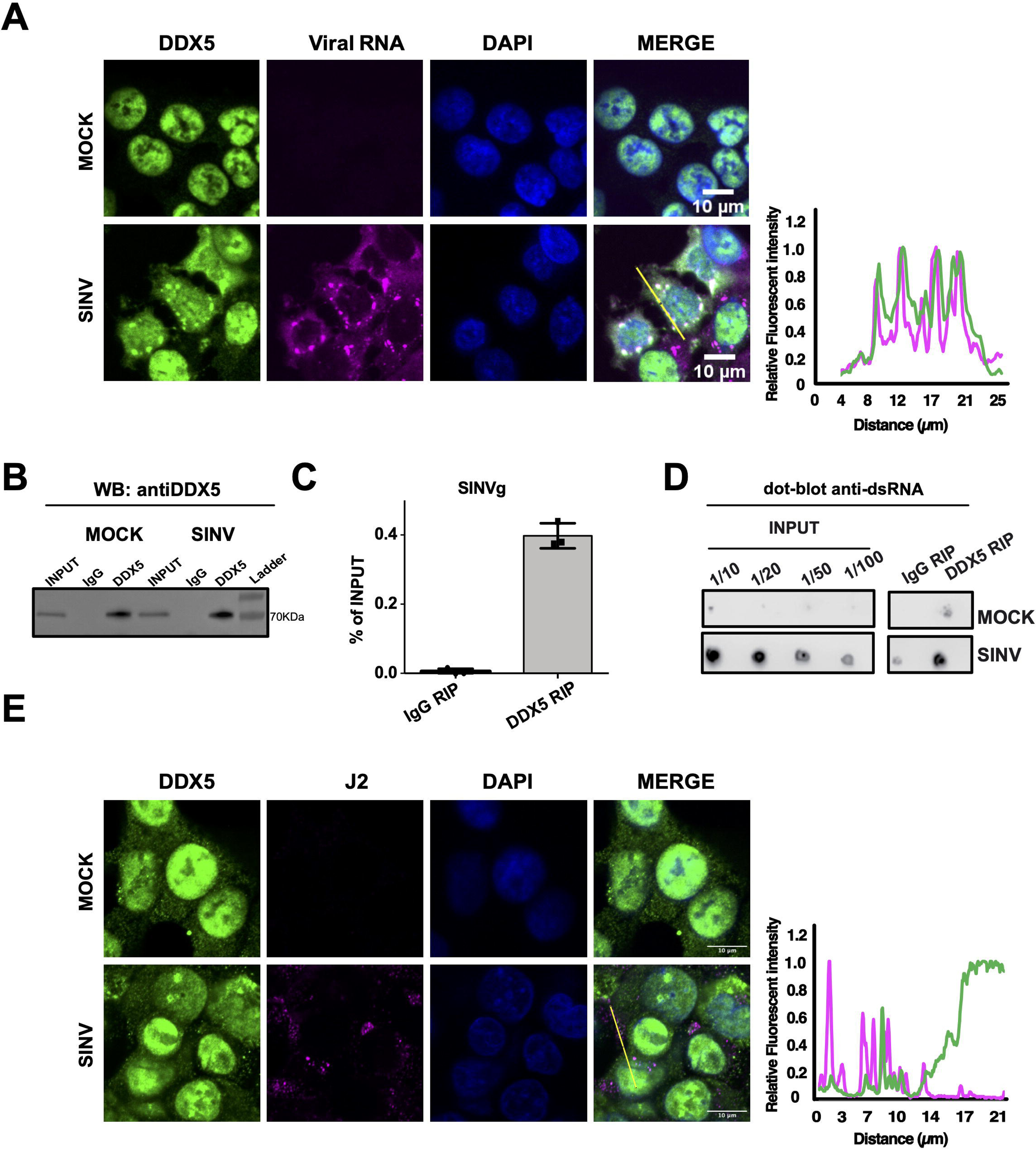
DDX5 interacts with the viral RNA. **A**) Confocal microscopy analysis of SINV (+) RNA and DDX5 protein localization in mock and infected HCT116 cells at 24 hpi by RNA fluorescence in situ hybridization (FISH) (in magenta) combined with protein immunostaining (in green). DAPI staining (in blue) and merge of the different channels are shown. Scale bar, 10µm. Representative fluorescence intensity profiles of DDX5 (green curve) and SINV RNA (magenta curve) along the yellow line represented on the merge panel and normalized to the maximum value are shown on the right. **B**) Anti DDX5 western blot on total lysate (INPUT), IgG-IP or DDX5-IP samples in both mock and SINV-GFP infected HCT116 cells. **C**) RT-qPCR on SINV genomic (g) RNA upon DDX5 RIP or IgG RIP. Results are expressed as percentage of Input (total RNA) and represent the mean ± standard deviation (SD) of three biological replicates (n = 3). **D**) Anti-dsRNA dot blot assay on serial dilutions of the total RNA (INPUT) and the undiluted RNA samples from DDX5-RIP or IgG-RIP, in mock and SINV infected conditions. J2 antibody was used to detect dsRNAs. **E**) Confocal co-immunofluorescence analysis was performed in mock and SINV infected HCT116 cells using anti-DDX5 rabbit antibody (in green) or anti-J2 mouse antibody (in magenta). Nuclei were stained with DAPI (in blue). Scale bar 10µm. Representative fluorescence intensity profiles of DDX5 (green curve) and J2 (magenta curve) along the yellow line represented on the merge panel and normalized to the maximum value are shown on the right.

Overall, these results suggest that DDX5 can bind to viral RNAs in the cytoplasm of SINV infected cells and localizes in close proximity to cytoplasmic viral dsRNA structures.

### 3.2. DDX5 interacts with host and viral proteins in SINV-infected conditions

Once we determined that DDX5 could be recruited to the viral RNA, we aimed to test whether any interaction between DDX5 and the viral proteins could also occur to modulate this association upon SINV infection. We immunoprecipitated DDX5 from mock and SINV-infected HCT116 cells at MOI 0.1 for 24 hours and eluates were analysed by liquid chromatography coupled to tandem mass spectrometry (LC-MS/MS). An IgG antibody was used as a negative control. Significantly enriched proteins were determined as having an FDR<1%, at least 5 spectral counts in the most abundant condition, a fold change of at least 2 and an adjusted p-value lower than 0.05.

Differential expression analysis revealed a total of 206 human proteins enriched in uninfected DDX5-IP over IgG-IP conditions (**Figure 2A** and **Table S1**), while 177 proteins were enriched in DDX5 IP over IgG IP control in SINV-infected samples (**Figure 2B and Table S2**).

**Figure 2:**
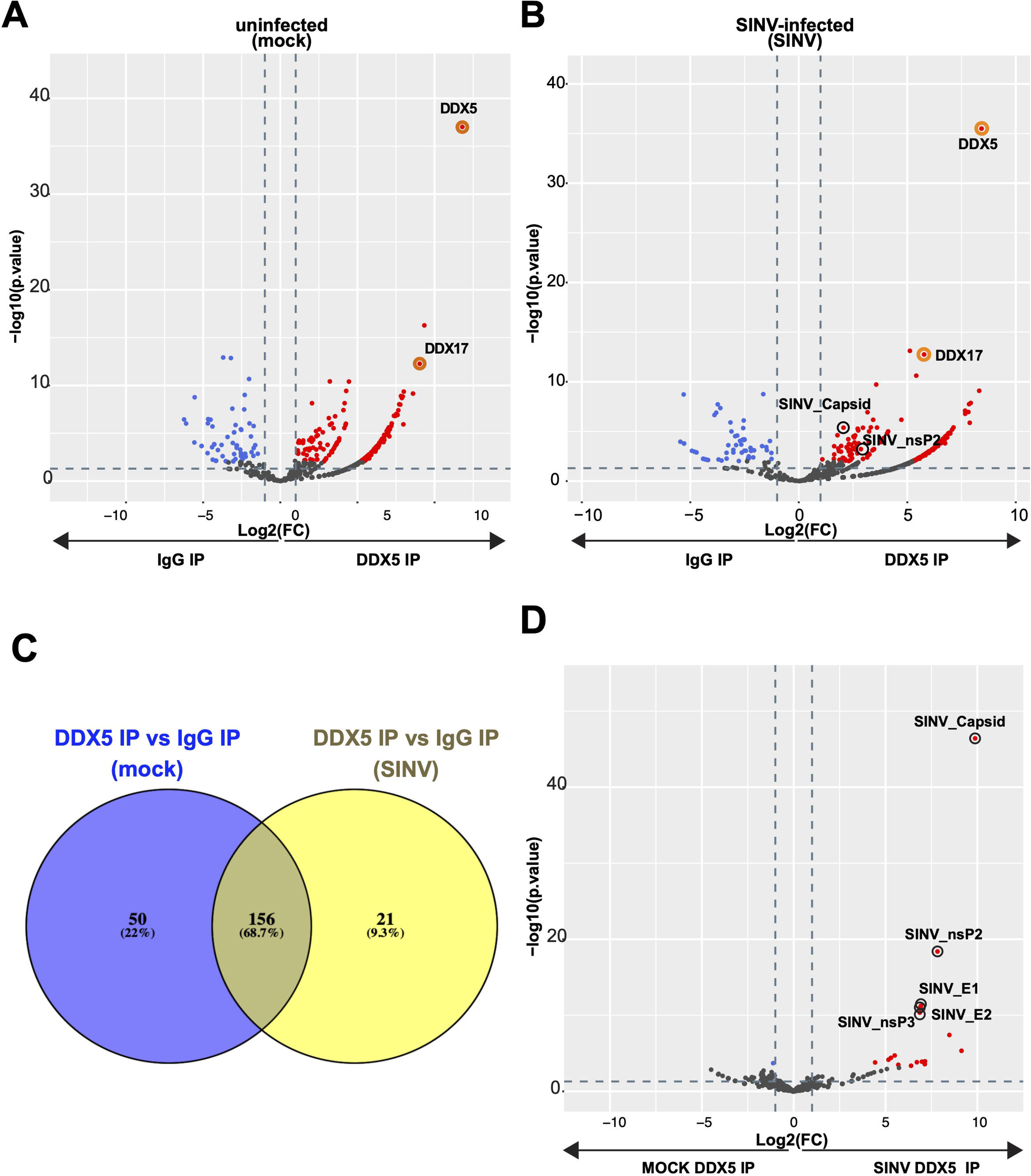
DDX5 interactome in mock and SINV-infected cells. **A, B and D)** Volcano plots showing the global protein enrichment in (**A**) mock-DDX5-IP versus mock-IgG-IP, (**B**) SINV-DDX5-IP versus mock-DDX5-IP and (**D**) SINV-DDX5-IP versus SINV-IgG-IP using data from three independent replicates. The cells were infected with SINV-GFP during for 24h at an MOI 0.1. Significantly enriched or depleted proteins are represented in red or blue, respectively. Cut off values: FDR<1%, at least 5 spectral counts in the most abundant condition, fold change > 2, adjusted p-value lower than 0.05. Viral proteins and selected host proteins are circled in black and orange, respectively. **C**) Venn diagram showing the number of overlapping and unique significantly enriched proteins identified in DDX5 IP compared to IgG in mock and SINV infected proteomic analyses.

Comparison of the DDX5 enrichment profiles in mock and in SINV conditions showed that 156 human proteins overlapped in the mock and SINV DDX5 interactomes, suggesting an overall stable DDX5 interactome core before and after infection (**Figure 2C**). Among them, we identified the polyribonucleotide nucleotidyltransferase 1 (PNPT1), an enzyme predominantly localized in the mitochondrial intermembrane space which acts as a 3’-to-5’ exoribonuclease and regulates mitochondrial RNA metabolism thereby modulating innate immunity (35). Even though PNPT1 and DDX5 were both enriched in our previous DRIMS experiments upon SINV infection (26), their close interaction has not been reported so far, Several RNA helicases such as DHX9, DHX30, DHX15, DDX3X, DDX21, DDX24, DDX54, DDX18, DDX10, DDX52 and DDX56, were also found in the DDX5 interactome. In particular, one of the most enriched human proteins in both mock and SINV-infected DDX5 IP was its well-known interactor DDX17 (10).

The viral capsid and nsP2 proteins were the most significantly enriched proteins in DDX5 IP SINV over DDX5 IP mock (**Figure 2D** and **Table S3**) and the only viral proteins significantly identified in DDX5 IP over IgG IP upon SINV infection (**Figure 2B and Table S2).** We thus decided to assess whether these association between DDX5 and these SINV proteins was specific and whether it relied on the binding to the viral RNA.

### 3.3. DDX5 interaction with the viral capsid is RNA-independent

To further characterise the interaction between DDX5 and the viral proteins, we chose to test all four SINV non-structural proteins (nsP1, nsP2, nsP3, nsP4) required for viral RNA synthesis and the structural capsid protein (CP) which associates with the viral genomic RNA for its packaging (**Figure 3A**). We individually cloned each of their coding sequence in order to express them with an N-terminal Flag-HA tag and verified their expression upon transient transfection in HCT116 cells by western blot analysis using an anti-Flag antibody (**Figure 3B, upper panel**). We next investigated whether the endogenous DDX5 co-immunoprecipitated with any of the overexpressed viral proteins in uninfected HCT116 cells in the absence of viral RNA or any other viral protein. While we did not confirm the interaction between nsP2 and DDX5 in the absence of viral RNA, we observed that the capsid protein associated with DDX5 independently of the viral RNA (**Figure 3B, lower panel**).

**Figure 3:**
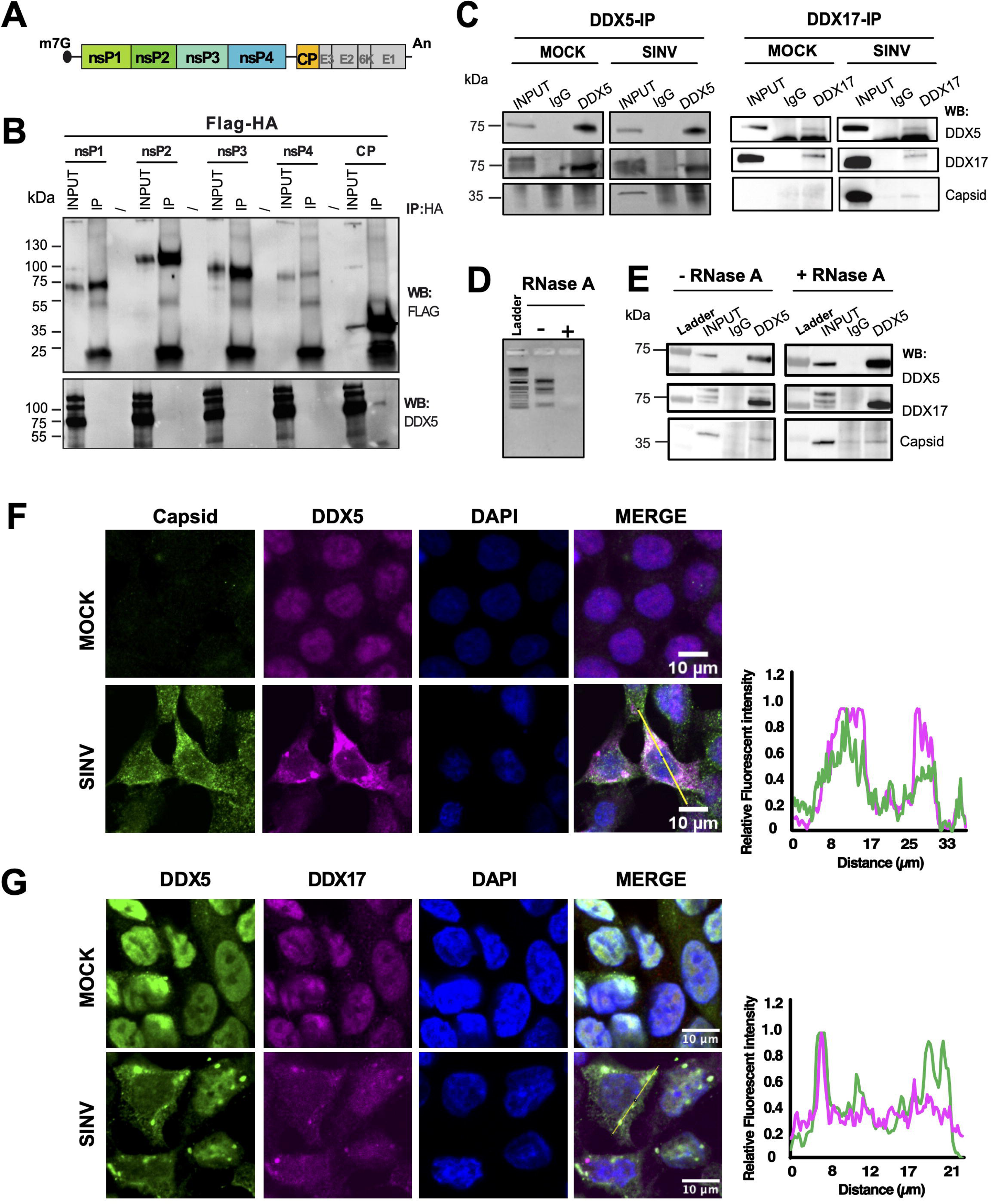
DDX5 interacts with the SINV capsid protein. **A**) Schematic illustration of SINV genomic organization. The 5’ cap and poly(A) tail are depicted. The non-structural proteins (np1, nsp2, nsp3, nsp4) from the first ORF (in green) and the structural proteins (capsid, E3, E2, 6K, E1) from the second ORF (in yellow and grey) are shown. The sequences coding for nsp1, nsP2, nsP3, nsP4 and capsid proteins were individually cloned and fusd with an N-terminal FLAG-HA tag. **B**) Western blot on total lysate (INP) or anti-HA IP (IP) from transiently transfected HCT116 cells using anti-FLAG and anti-DDX5 antibodies. Position and molecular weight (KDa) of protein size markers are shown on the left. **C)** Western blot on total lysate (INPUT), IgG-IP (IgG), DDX5-IP (DDX5) or DDX17-IP (DDX17) samples in both mock and SINV-GFP infected lysates, using antibodies against DDX5, DDX17 and viral capsid. **D)** Representative agarose gel showing the total RNA before and after RNase treatment. **E)** Western blot on total lysate (INPUT), IgG-IP (IgG) or DDX5-IP (DDX5) samples in both mock and SINV-GFP infected lysates, treated or not with RNase A, using antibodies against DDX5, DDX17 and viral capsid. **F-G**) Confocal co-immunofluorescence analysis performed in mock and SINV-infected HCT116 cells using specific primary antibodies. Rabbit anti-capsid (in green) and mouse anti-DDX5 antibody (in magenta) are shown in (**F**). Rabbit anti-DDX5(in green) and mouse anti-DDX17 antibodies (in magenta) are shown in (**G**). DAPI was used to stain the nuclei (in blue). Scale bar 10µm. Representative fluorescence intensity profiles of capsid (green curve) and DDX5 (magenta curve) **(F)** or DDX5 (green curve) and DDX17 (magenta curve) **G**) along the yellow line represented on the merge panel and normalized to the maximum value are shown on the right.

We further validated the interaction of DDX5 and the viral capsid by co-immunoprecipitation experiments upon SINV infection (**Figure 3C, left panels**). Reciprocal co-immunoprecipitation of DDX17 confirmed its interaction with DDX5 and the viral capsid (**Figure 3C, right panels**), suggesting that the three proteins may be in close proximity in infected conditions. Of note, the interaction between DDX5 and capsid as well as the association with DDX17 are mostly maintained upon RNase A treatment. (**Figure 3D-E**).

We then performed co-immunostaining analyses to verify whether DDX5 co-localizes with the viral capsid in SINV-infected HCT116 cells. Confocal microscopy analysis demonstrated that endogenous DDX5 co-localizes with the capsid protein in SINV-infected cells supporting the possibility that the two proteins interact upon infection (**Figure 3F**).

Interestingly, while DDX5 and its cofactor DDX17 colocalize in the nucleus in uninfected conditions, both proteins re-localize to the cytoplasm in discrete perinuclear foci upon SINV infection (**Figure 3G**) most likely in proximity of viral replication sites (see **Figure 1A**).

Overall, these results confirm the interaction between the viral capsid and DDX5 in infected cells and indicate that DDX17 interaction with DDX5 and capsid can occur in the cytoplasm of the infected cells.

### 3.4. DDX5 depletion reduces SINV infection in HCT116 cells

To formally determine the impact of DDX5 on SINV replication and infection, we first knocked it down by siRNA treatment in HCT116 cells, which were then infected for 24h with SINV-GFP at an MOI of 0.1. We observed a reduction in viral capsid protein level by western blot upon DDX5 depletion (siDDX5) compared to the control condition (siCTRL) (**Figure 4A**). In addition, we found that the viral genomic (g)RNA level (**Figure 4B**) and the viral titer (**Figure 4C**) were also reduced in siDDX5 compared to siCTRL-treated cells.

**Figure 4:**
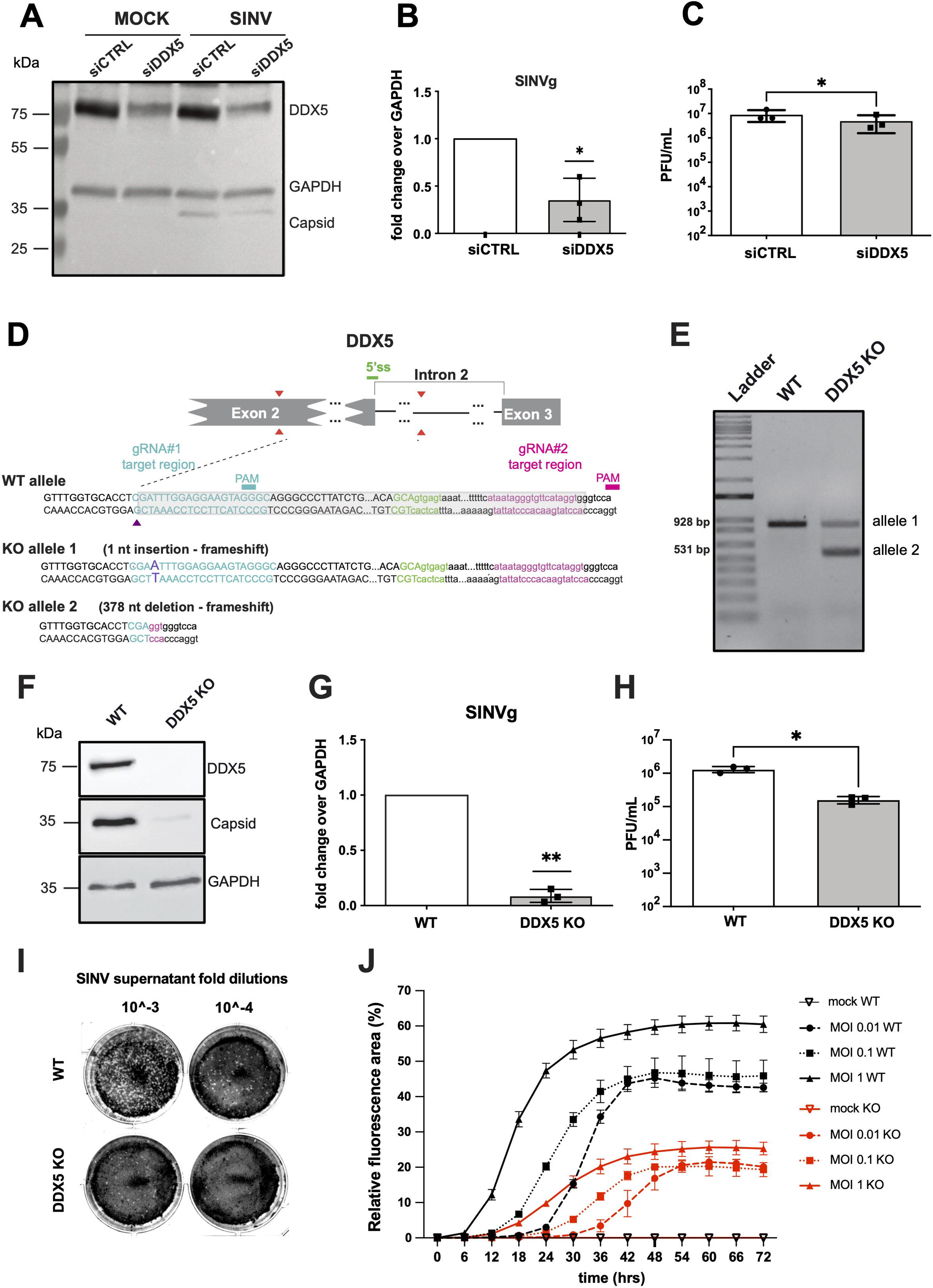
SINV infection is reduced in DDX5*-*depleted HCT116 cells. **A**) Western blot on HCT116 mock or SINV-GFP infected cells upon siCTRL and siDDX5 transfection. DDX5 and viral capsid were detected by specific primary antibodies. GAPDH was used as loading control. Position and molecular weight (KDa) of protein size markers are shown on the left. **B**) RT-qPCR of SINV genomic (g) RNA in SINV-GFP infected HCT116 cells (MOI 0.1, 24 hpi), upon siDDX5 and siCTRL transfection. Results are expressed as relative to GAPDH and normalized to the siCTRL condition. **C**) SINV-GFP viral titers from infected HCT116 cells transfected with siCTRL or siDDX5 was quantified by plaque assay. Results in **B-C** represent the mean ± standard deviation (SD) of three biological replicates (n = 3). * p ≤ 0.05, (**B**) one-sample t test or (**C**) paired Student’s t test. **D**) Schematic diagram of the *DDX5* CRIPR/Cas9 KO strategy. Two gRNAs targeting *DDX5* exon 2 and intron 2 respectively were used to delete the 5’ splice site (ss) region (5’ss is shown in green, #1 target region in blue and #2 target region in magenta). The purple arrow represents an indel in *DDX5* exon 2 for the KO allele 1. The grey rectangle represents a 378 nucleotide (nt) deletion in KO allele 2. **E**) Agarose gel showing the PCR fragments amplified from cellular genomic DNA, corresponding to the WT and KO alleles, respectively, which were gel purified and sequenced. Amplicon size is shown: the upper band corresponds to KO allele 1, the lower band corresponds to the KO allele 2. **F)** Western blot on mock or SINV-GFP infected cells in WT or *DDX5* KO HCT116 cells. Antibodies against DDX5 and viral capsid were used. GAPDH was used as loading normalizer. **G**) Level of SINV genomic RNA relative to GAPDH monitored by RT-qPCR in WT or *DDX5* KO HCT116 cells upon SINV-GFP infection at MOI 0.1 for 24 hours. Results are plotted as fold change relative to WT cells. **H**) Viral particles of SINV-GFP from the supernatant of SINV-GFP infected WT or *DDX5* KO HCT116 cells were quantified by plaque assay on Vero cells by plaque assay. The results from three independent experiments are presented here. **I**) Representative plaque assay images after crystal violet staining of Vero E6 cells infected with SINV-GFP supernatant at the indicates dilutions (10^-3^ and 10^-4^) from infected WT or *DDX5* KO HCT116 cells. **J**) SINV-GFP infection kinetics in WT and *DDX5* KO HCT116 cells. The relative GFP fluorescence area (expressed in percentage) of WT and *DDX5* KO HCT116 cells as a function of time was measured after SINV GFP infection at an MOI of 0.01 (left panel), 0.1 (middle panel) and 1 (right panel) every 6 hours for 72 hours with a CellcyteX automated cell counter and analyser. WT HCT116 cells, black; *DDX5* KO HCT116 cells, red. Error bars in **G-H** and **I** represent the mean ± standard deviation (SD) of three independent biological experiments. * p ≤ 0.05, **p ≤ 0.01, (**G**) one-sample t test or (**H**) paired Student’s t test.

To further establish the negative effect of DDX5 depletion on SINV infection, we generated homozygous *DDX5* knockout (KO) HCT116 cells by CRISPR/Cas9 (**Figure 4D**). The *DDX5* KO clone was characterized at the genomic DNA level (**Figure 4E**) and the absence of DDX5 protein was verified by western blot (**Figure 4F**).

Upon SINV-GFP infection for 24h at an MOI of 0.1 we observed a strong reduction in viral capsid protein in *DDX5* KO HCT116 compared to WT HCT116 cells by western blot (**Figure 4F**). The viral genomic RNA level (**Figure 4G**) and the viral titer (**Figure 4H-I**) were also significantly reduced in the absence of DDX5.

We then measured the relative fluorescence of GFP to quantify the impact of *DDX5* knockout during a time course of SINV infection using a CellcyteX automated cell counter and analyser. A robust decrease in GFP fluorescence signal was observed in SINV-GFP infected *DDX5* KO compared to WT HCT116 cells at three MOIs tested (MOI of 0.01, 0.1 and 1) over 72 hours (**Figure 4J**). These results indicate that DDX5 depletion strongly reduces the infection over time and independently on the MOI used already at the earliest step of the viral replication cycle. Taken together our results point to a negative impact of DDX5 depletion on SINV infection in our experimental conditions suggesting that DDX5 is a pro-viral host factor for SINV infection.

### 3.5. DDX5 re-expression increases capsid levels in SINV-infected HCT116 cells

To better assess the pro-viral function of DDX5 on SINV, we stably re-expressed a V5-DDX5 or BFP protein (control) in *DDX5* KO HCT116 cells and verified that a partial DDX5 expression could be restored in our experimental conditions compare to WT HCT116 cells (**Figure 5A**). We then analysed the effect of the ectopic DDX5 expression on the virus at 24hpi at an MOI of 0.1 and observed an increase in viral capsid protein level by western blot (**Figure 5B**) in V5-DDX5 cells compared to the control BFP one infected with SINV-GFP. In contrast, we found that the viral genomic (g)RNA, sub-genomic (sg)RNA level (**Figure 5C-D**) and the viral titer (**Figure 5E**) were not significantly affected in V5-DDX5 compared to BFP expressing HCT116 KO cells at this specific time point.

**Figure 5:**
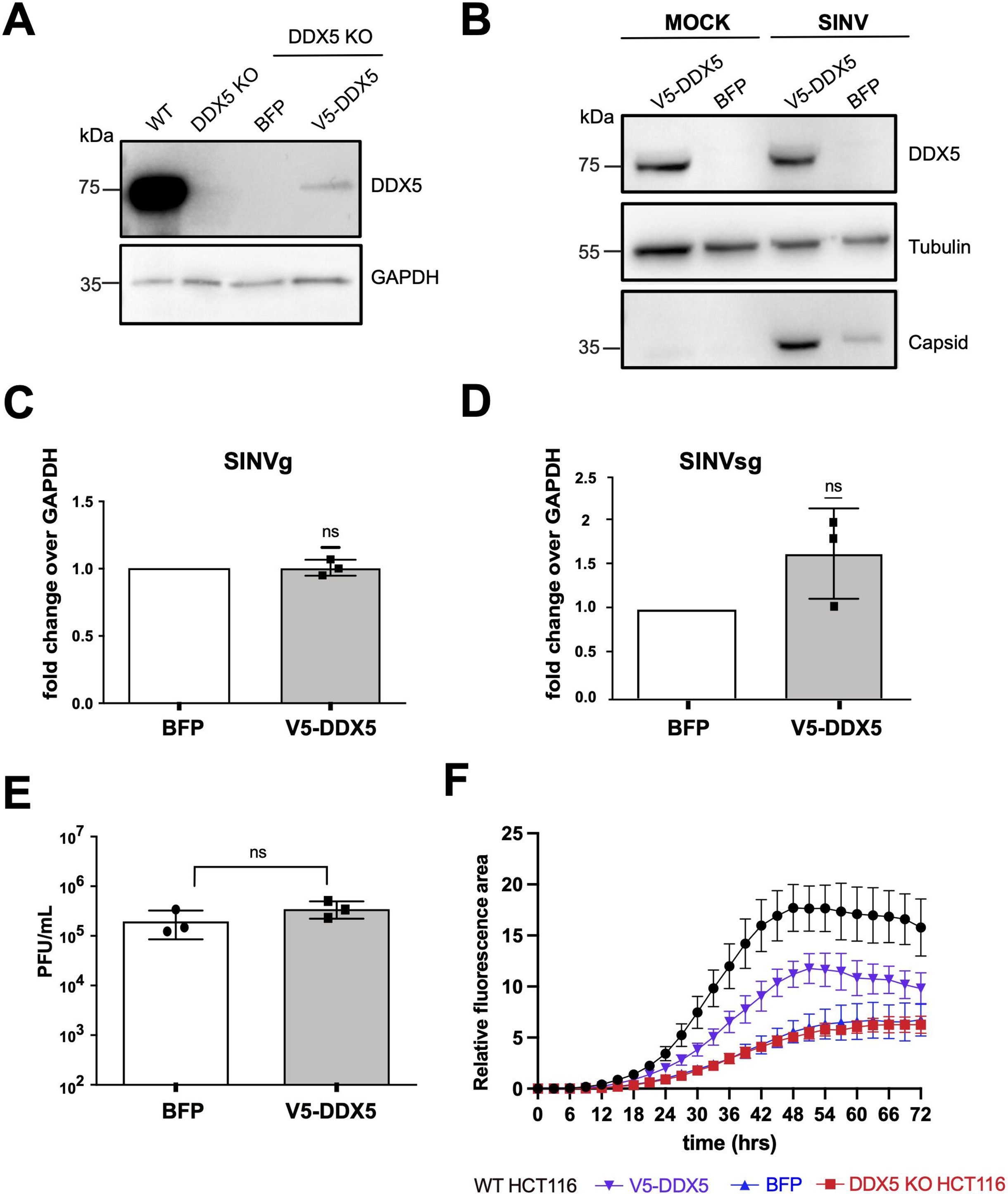
Effect of DDX5 re-expression on SINV in *DDX5* KO HCT116 cells. **A**) Western blot on WT, *DDX5* KO HCT116 and *DDX5* KO HCT116 stably expressing either V5-DDX5 (V5-DDX5) or BFP (BFP). Antibodies directed against DDX5 and GAPDH antibody were used. **B)** Western blot on SINV-GFP infected *DDX5* KO HCT116 overexpressing V5-DDX5 or BFP (24 hpi, MOI 0.1). Antibodies directed against DDX5 and capsid were used. Tubulin antibody was used as loading control. (**C-D**) RT-qPCR of SINV genomic (g) RNA (**C**) and SINV sub-genomic (sg) RNA (**D**) in SINV-GFP infected *DDX5* KO HCT116 stably expressing V5-DDX5 or BFP (MOI 0.1, 24 hpi). Results are expressed as relative to GAPDH and normalized over the siCTRL condition. **E**) SINV-GFP viral titers from SINV-GFP infected *DDX5* KO HCT116 stably expressing V5-DDX5 or BFP (MOI 0.1, 24 hpi), quantified by plaque assay. Results in (**C-D-E)** represent the mean ± standard deviation of three biological replicates (n = 3). ns = non significant, (**C-D**) one sample t test or (**E**) paired Student’s t test. **F)** SINV-GFP infection kinetics in WT, *DDX5* KO HCT116 and *DDX5* KO HCT116 stably expressing either V5-DDX5 (V5-DDX5) or BFP (BFP). The relative GFP fluorescence area (expressed in percentage) of the infected cells as a function of time was measured after SINV GFP infection at an MOI of 0.1, every 3 hours for 72 hours with a CellcyteX automated cell counter and analyser. WT HCT116 cells, black; *DDX5* KO HCT116 cells, red; V5-DDX5 cells, purple; BFP cells, blue.

Finally, while the effect was not as pronounced as in WT cells, the quantification of the relative GFP fluorescence during a time course of SINV-GFP infection (MOI 0.1) revealed a partial rescue of the GFP signal in the V5-DDX5 cells compared to the BFP and the *DDX5* KO ones over time (**Figure 5F**).

Taken together, these observations suggest that the ectopic DDX5 expression has a positive impact on SINV-GFP infection indicating an overall pro-viral effect on the virus. However, the impact of DDX5 overexpression on the virus at 24hpi is mostly observed as an increase of the viral capsid protein level and does not induce a significant increase on viral RNA and infectious particle production, although we could observe a tendency for an increase in the presence of DDX5. Nonetheless, the kinetic of GFP fluorescence measurement clearly indicates that the phenotype of the DDX5 knock-out can be complemented by its re-expression.

### 3.6. Combined DDX5 and DDX17 depletion has a synergic detrimental impact on SINV

Once established that DDX5 depletion has a negative impact on SINV and that its re-expression in a *DDX5* KO background has a positive impact on the virus, we set to determine whether DDX17 plays a role together with or independently of DDX5 during SINV infection. We first knocked down DDX17, then infected HCT116 cells with SINV-GFP. We validated the knockdown efficiency by western blot compared to an siRNA control and observed that viral capsid levels were reduced upon siDDX17 treatment (**Figure 6A**). In addition, SINV genomic and sub-genomic RNA levels (**Figure 6B-C**) as well as SINV viral particle production (**Figure 6D**) were also significantly reduced in DDX17 knockdown conditions compared to siCTRL. These results suggested that just like DDX5, DDX17 has a positive effect on SINV infection.

**Figure 6:**
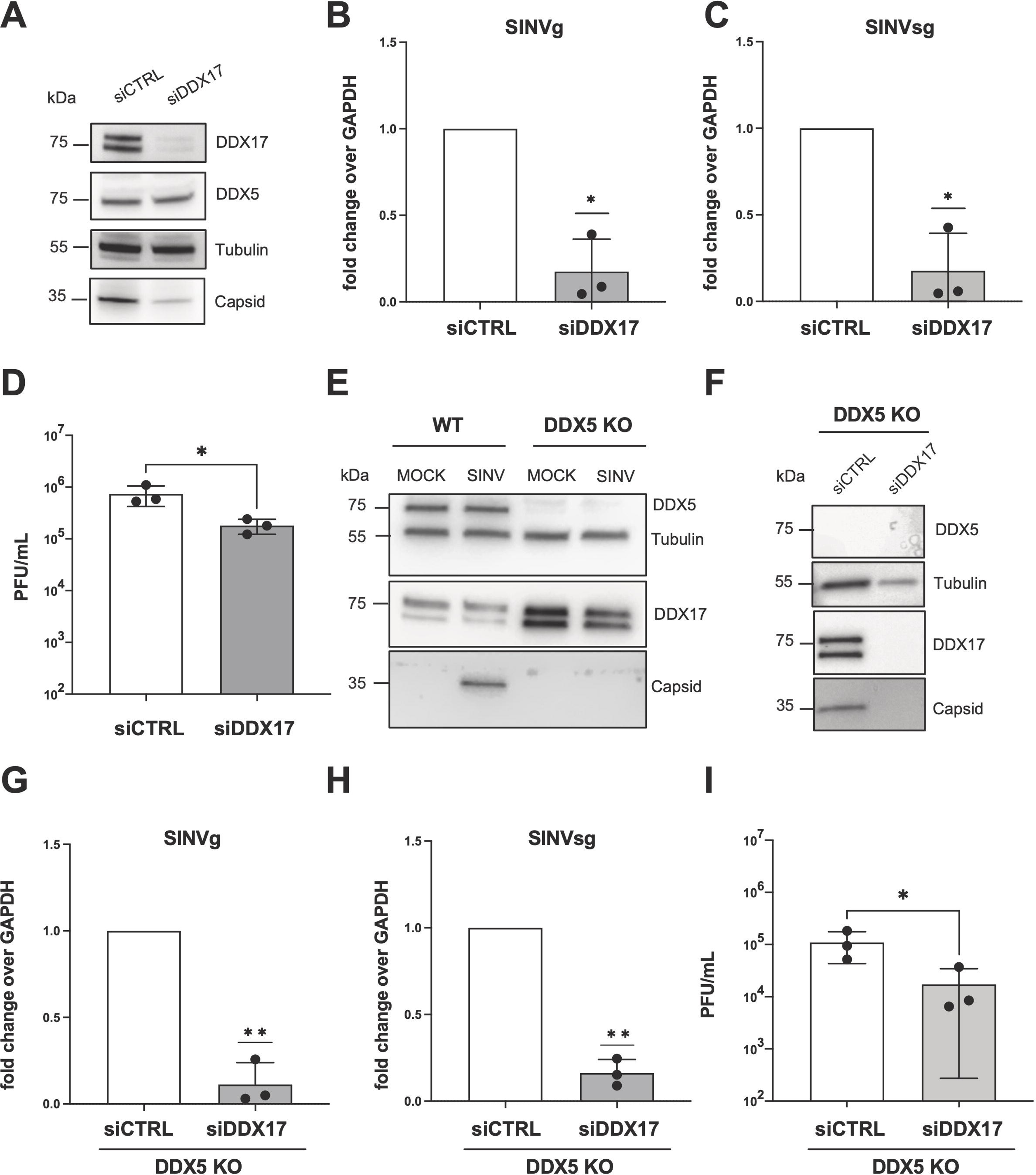
Effect of DDX17 depletion on SINV in WT and *DDX5* KO HCT116 cells. **A**) Western blot on SINV-GFP infected HCT116 cells, treated with siCTRL and siDDX17. Antibodies directed against DDX5, DDX17 and capsid were used. Tubulin antibody was used as loading control. (**B-C**) RT-qPCR of SINV genomic (g) RNA (**B**) and SINV sub-genomic (sg) RNA (**C**) in siRNA treated SINV-GFP infected HCT116 cells (MOI 0.1, 24 hpi). Results are expressed as relative to GAPDH and normalized over the siCTRL condition. **D**) SINV-GFP viral titers from infected HCT116 cells transfected with the indicated siRNAs, quantified by plaque assay. **E)** Western blot on mock SINV-GFP infected WT and *DDX5* KO HCT116 cells. Antibodies directed against DDX5, DDX17 and capsid were used. Tubulin antibody was used as loading control. **F)** Western blot of samples from SINV-GFP infected *DDX5* KO HCT116 cells upon siCTRL or siDDX17 transfection. Specific primary antibodies against DDX5, DDX17 and viral capsid were used. Tubulin was used as loading control. Position and molecular weight (KDa) of protein size markers are shown on the left. **G-H)** RT-qPCR of SINV genomic (g) RNA (**G**) and SINV sub-genomic (sg) RNA (**H**) in SINV-GFP infected *DDX5* KO HCT116 cells (MOI 0.1, 24 hpi), upon siRNA transfection. Results are expressed as relative to GAPDH and normalized to the siCTRL condition. **I)** SINV-GFP viral titers from infected *DDX5* KO HCT116 cells transfected with the indicated siRNAs, quantified by plaque assay. Results in (**B-C-D** and **G-H-I)** represent the mean ± standard deviation of three biological replicates (n = 3). * p ≤ 0.05, **p ≤ 0.01, (**B-C** and **G-H**) one sample t test or (**D** and **I**) paired Student’s t test.

We proceeded to investigate whether DDX17 acts in synergy or competition with DDX5 by analysing the impact of DDX17 knockdown in *DDX5* KO HCT116 cells where viral infection is already weakened (see **Figure 4D-J**). As previously reported (13), the protein levels of both DDX7 isoforms (p72 and p82) increased in *DDX5* KO HCT116 compared to WT HCT116 cells and such an effect was independent on the infection (**Figure 6E**). However, upon SINV-GFP infection, we observed a further reduction of the viral capsid expression in siDDX17 treated *DDX5* KO HCT116 cells compared to the control condition by western blot (**Figure 6F**). SINV genomic and sub-genomic RNA levels (**Figure 6G-H**), as well as SINV viral particle production (**Figure 6I**) were significantly diminished in DDX17 knockdown conditions compared to siCTRL in *DDX5* KO HCT116 cells. Overall, these results indicate that DDX5 and DDX17 may have independent pro-viral functions during SINV infection because their individual depletion causes reduction of viral infection at the protein, RNA and viral production levels and their joint depletion has a cumulative negative effect on SINV infection.

## 4. Discussion

The human DEAD-box RNA helicase DDX5 plays a pivotal role in virus–host interaction being mostly pro-viral for several viral families. In this study, we investigated and extended the function of DDX5 as a cellular cofactor of SINV replication in HCT116 cells. Despite being an uncommon model to study alphavirus infection, the HCT116 cell line was chosen because it can be easily infected with SINV and can be used for CRISPR-Cas9 gene editing procedures (36).

We found that DDX5 interacts with viral RNA at the cytoplasmic viral replication sites, which may be important for its pro-viral activity. RNA helicases recognize the RNA structure rather than their sequence and they have shown little substrate specificity *in vitro* (1). This lack of specificity is counterbalanced by their association with specific interactors *in vivo*. For instance, DDX5 interaction with either viral non-structural or structural proteins is essential for its pro-viral role across different RNA virus infections (14).

Our proteomic analysis revealed the interaction between DDX5 and SINV capsid, as well as with the nsP2 protein. The cytoplasmic SINV capsid protein is essential for genomic RNA packaging and the first steps of viral morphogenesis. In contrast, SINV nsP2 is the protease and helicase of the viral replication complex and it is the only viral protein known to re-localise to the nucleus to inhibit host transcription (37). The interaction between DDX5 and nsP2 may thus occur both in the nucleus and in the cytoplasm of the infected cell. The association between the viral capsid and DDX5 was further supported by co-immunoprecipitation experiments and imaging analyses. This interaction is RNA-independent since it occurs both in uninfected cells expressing a tagged version of the capsid protein and in the presence of an RNase treatment in SINV infected cells.

We showed that DDX5 depletion has a detrimental effect on viral replication and production in infected cells. Furthermore, we established a stable cell line that ectopically expressed DDX5 in a *DDX5* KO background to assess its functional impact on the virus. Despite its modest overexpression, the presence of DDX5 was sufficient to partially rescue the GFP levels in a time course of SINV-GFP infection. Although we did not observe a significant increase of viral gRNA, sgRNA or titers, we noted an enhanced accumulation of the capsid protein in DDX5 KO cells upon re-expression of a tagged DDX5. This partial effect might depend on the relatively low expression levels of the V5-DDX5 or on the choice of the inappropriate time point of infection.

Indeed, we observed a partial rescue of fluorescence in cells expressing V5-DDX5 compared to the controls when we monitored GFP expression from the virus in real time over 72 hours. Taken together, these results support the pro-viral role of DDX5 on SINV.

The DDX5 interactome analysis also revealed associations with a number of cellular RNA binding proteins that could have a function during SINV infection. These include several RNA helicases of the DEAD/H family and other interesting factors such as PNPT1. Interestingly, we previously showed that DDX5, DDX17 and PNPT1 are an integral part of the SINV dsRNA associated proteome (26). Further investigations are needed to better characterize if the association of those novel DDX5 interactors have functional consequences on SINV infection. In this study we focused on DDX17 which was previously known to associate with DDX5.

We demonstrated by co-immunoprecipitation experiments that DDX17 interacts with DDX5 and the viral capsid in infected cells. It is now established that the replication of positive strand RNA viruses in the cytoplasm can induce re-localization of nuclear host proteins and promote infection (38). Of note, we showed that DDX5 re-localizes from the nucleus to the cytoplasm with DDX17 upon SINV infection where they can both interact with the viral capsid protein. It is thus conceivable that the cofactor DDX17 operates with DDX5 within the same complex, carrying out a synergistic or additive function during infection. A previous study showed that Rm62, the fly orthologue of DDX17 and DDX5, did not affect SINV infection in *D. melanogaster*, and that DDX17 knockdown showed no antiviral effect on the accumulation of SINV capsid protein in human U2OS cells (16). In contrast, we demonstrated that the single depletion of DDX17 reduces SINV infection in HCT116 cells and its combined depletion with DDX5 in the KO background further decreases viral levels, suggesting a synergism between the two proteins in promoting viral infection.

These experiments also ruled out the possibility that the observed reduction in SINV replication upon DDX5 depletion was due to an antiviral effect of compensatory DDX17 overexpression.

As both DDX5 and DDX17 inhibition reduces virus levels, further investigation is required to better characterize these factors as potential targets for antiviral therapies. It will also be important to expand our functional observations to other cellular models more relevant for alphavirus infection in the future.

Additional analyses will be needed to formally assess whether DDX5 direct action supports viral replication and if its binding to viral RNA and helicase activity is necessary for its function. Being involved in many aspects of cellular RNA biology, one cannot formally exclude the contribution of DDX5 indirect effects on infection. For instance, DDX5 activity as a regulator of N6-methyladenosine levels on the DHX58 and NF-κB mRNAs could negatively impact antiviral innate immunity and therefore enhance viral infection (39). Finally, it could be of interest to determine whether the helicase activity of DDX5 is required for its pro-viral effect on SINV production and which protein domains are important for the interaction with the capsid viral protein.

## 5. Conclusion

In conclusion, this study uncovers DDX5 as a novel positive host factor for the prototype alphavirus SINV and significantly advances our understanding of the role played by RNA helicases in viral infections. Our results demonstrate the interaction of DDX5 with viral RNA, suggesting a potential impact on early stages of viral replication. We discovered a re-localization of DDX5 together with its partner DDX17 to the cytoplasm upon SINV infection and their association with the viral capsid protein. Depletion of both DDX5 and DDX17 strongly impacts SINV infection. Of note, the detrimental effect caused by DDX5 depletion on the virus is partially offset by DDX5 ectopic expression, providing evidence for the pro-viral function of DDX5 on SINV.

## Funding

This work received financial support from the Agence Nationale de la Recherche (EndoDRAI project ANR-22-CE15-0011-01 (to E.G.) and was supported by the Idex 2022 Attractivité grant under the framework of the IdeX University of Strasbourg (to E.G.) and from the French Minister for Higher Education, Research and Innovation (PhD contract to C.P.). This work of the Interdisciplinary Thematic Institute IMCbio+, as part of the ITI 2021-2028 program of the University of Strasbourg, CNRS and Inserm, was supported by IdEx Unistra (ANR-10-IDEX-0002), by SFRI-STRAT’US project (ANR-20-SFRI-0012), EUR IMCBio (IMCBio ANR-17-EURE-0023) under the framework of the French Investments for the Future Program. S.P. also receive funding from ANR (Project RIFLAVIRAM ANR-21-CE35-0018-01). The mass spectrometry instrumentations were funded by the University of Strasbourg, IdEx “Equipement mi-lourd” 2015.

## Contributions

**MM**: Conceptualization, Methodology, Validation, Investigation, Visualization, Writing - Original Draft. **LP**: Investigation. **CP**: Investigation. **ND**: Investigation. **JC**: Investigation, Validation. **PH**: Investigation, Validation. **SP**: Conceptualization., Writing - Review & Editing, Funding acquisition. **EG**: Conceptualization, Methodology, Validation, Investigation, Visualization, Writing - Review & Editing, Funding acquisition, Supervision.

## Acknowledgements

We thank Béatrice Chane-Woon-Ming for her help with the proteomic analyses, Dr. Paula Lopez for her initial support with plaque assays and Agathe Hunckler for technical assistance. We also thank Dr. Carla Saleh for providing the SINV WT and GFP infectious clones and Dr. Diane Griffin for providing the CAPSID antibody. We thank Dr. Cyril Bourgeois, Dr. Richard Patryk Ngondo and members of the Pfeffer labs for critical reading of the manuscript.

## Ethics declarations

### Ethics approval and consent to participate

Not applicable.

### Consent for publication

Not applicable.

### Competing interests

The authors declare that there are no conflicts of interest.

### Availability of data and materials

The mass spectrometry proteomics datasets supporting the conclusions of this article are available and have been deposited to the ProteomeXchange Consortium via the PRIDE (40) partner repository with the dataset identifier PXD044867 and 10.6019/PXD044867.

## Notes

### Competing Interest Statement

The authors have declared no competing interest.

### Summary of Updates

-Addition of new data in Figure 3 and 5. - Rewriting of the results, methods and discussion section to take into accounts all the above changes.

