## Supplementary material for "DEAD box RNA helicase 5 is a new pro-viral host factor for Sindbis virus infection": Supplmentary tables

**Supplementary tables**

**Table S1: List of significantly enriched proteins in mock-DDX5_IP versus mock-IgG_IP**

**1st column**: Protein name

**2nd column**: Column named as the treatment level with the mean raw spectral counts observed for this condition

**3rd column**: Column named as the control level with the mean raw spectral counts observed for this condition

**deltaSC**: Difference between mean raw spectral counts

**lFC.Av**: Log fold change computed from the mean expression levels taking into account the given normalization factors.

**logFC**: Log fold change estimated by fitting the given GLM model. The reference level of the main factor is taken as control.

**LR**: The likelihood ratio for edgeR.

**p.val**: The unadjusted p-values obtained from the tests.

**adjp**: The multitest adjusted p-values with FDR control.

|  | mock_DDX5IP | mock_IgG_IP | deltaSC | lFC_Av | LogFC | LR | p_value | adjp | negLog_adjp |
| --- | --- | --- | --- | --- | --- | --- | --- | --- | --- |
| DDX5_HUMAN | 745 | 0 | 745 | Inf | 11.8 | 164.9 | 9.65E-38 | 1.23E-34 | 33.91116 |
| DDX21_HUMAN | 136 | 0 | 136 | Inf | 9.338 | 70.22 | 5.31E-17 | 3.37E-14 | 13.47185 |
| DDX17_HUMAN | 109.7 | 0 | 109.7 | Inf | 9.036 | 52.04 | 5.44E-13 | 1.39E-10 | 9.85855 |
| RL6_HUMAN | 85.7 | 0.7 | 85 | 3.904 | 4.45 | 43.63 | 3.96E-11 | 6.30E-09 | 8.20087 |
| YBOX1_HUMAN | 120.3 | 2.3 | 118 | 3.687 | 3.213 | 43.71 | 3.80E-11 | 6.30E-09 | 8.20087 |
| PABP1_HUMAN | 74.3 | 0.7 | 73.6 | 5.681 | 4.273 | 39.23 | 3.76E-10 | 5.32E-08 | 7.27433 |
| SRRM2_HUMAN | 54.3 | 0 | 54.3 | Inf | 8.005 | 38.91 | 4.43E-10 | 5.63E-08 | 7.24934 |
| PNPT1_HUMAN | 81.3 | 0 | 81.3 | Inf | 8.602 | 37.99 | 7.11E-10 | 8.22E-08 | 7.08523 |
| NOP56_HUMAN | 47.7 | 0 | 47.7 | Inf | 7.821 | 36.84 | 1.29E-09 | 1.26E-07 | 6.90066 |
| YBOX3_HUMAN | 51.3 | 0 | 51.3 | Inf | 7.937 | 36.6 | 1.45E-09 | 1.32E-07 | 6.87976 |
| RL35_HUMAN | 49.7 | 0 | 49.7 | Inf | 7.883 | 35.93 | 2.05E-09 | 1.63E-07 | 6.78888 |
| RS6_HUMAN | 68.7 | 0.7 | 68 | 5.578 | 4.17 | 33.48 | 7.18E-09 | 5.11E-07 | 6.29183 |
| RL4_HUMAN | 182.3 | 7.7 | 174.6 | 2.104 | 2.056 | 33.47 | 7.23E-09 | 5.11E-07 | 6.29183 |
| RL36_HUMAN | 44 | 0 | 44 | Inf | 7.71 | 32.97 | 9.35E-09 | 6.26E-07 | 6.20336 |
| RL13A_HUMAN | 46.3 | 0 | 46.3 | Inf | 7.778 | 32.83 | 1.01E-08 | 6.40E-07 | 6.19389 |
| SRRM1_HUMAN | 42 | 0 | 42 | Inf | 7.65 | 30.17 | 3.96E-08 | 2.10E-06 | 5.67778 |
| DHX9_HUMAN | 65 | 0.7 | 64.3 | 5.5 | 4.095 | 30.24 | 3.83E-08 | 2.10E-06 | 5.67778 |
| RL21_HUMAN | 38.3 | 0 | 38.3 | Inf | 7.517 | 28.44 | 9.66E-08 | 4.91E-06 | 5.30865 |
| RL1D1_HUMAN | 40.7 | 0 | 40.7 | Inf | 7.606 | 28.02 | 1.20E-07 | 5.88E-06 | 5.23047 |
| NAT10_HUMAN | 34.7 | 0 | 34.7 | Inf | 7.369 | 27.6 | 1.49E-07 | 7.03E-06 | 5.15304 |
| NOP2_HUMAN | 36.7 | 0 | 36.7 | Inf | 7.457 | 27.52 | 1.56E-07 | 7.07E-06 | 5.15064 |
| RL10A_HUMAN | 65.3 | 1 | 64.3 | 4.928 | 3.555 | 27.36 | 1.69E-07 | 7.42E-06 | 5.12977 |
| RL17_HUMAN | 65.3 | 1.3 | 64 | 4.503 | 3.146 | 26.5 | 2.63E-07 | 1.12E-05 | 4.95273 |
| DDX24_HUMAN | 31.3 | 0 | 31.3 | Inf | 7.225 | 24.04 | 9.46E-07 | 3.20E-05 | 4.49444 |
| HS90A_HUMAN | 91.7 | 1.7 | 90 | 4.641 | 3.337 | 24.04 | 9.44E-07 | 3.20E-05 | 4.49444 |
| TOP1_HUMAN | 38.7 | 0.3 | 38.4 | 5.735 | 4.251 | 23.92 | 1.00E-06 | 3.27E-05 | 4.48545 |
| KI67_HUMAN | 53.3 | 0 | 53.3 | Inf | 7.98 | 23.63 | 1.17E-06 | 3.72E-05 | 4.42981 |
| RL18A_HUMAN | 52.3 | 1 | 51.3 | 4.597 | 3.222 | 23.42 | 1.30E-06 | 4.03E-05 | 4.39459 |
| RS3_HUMAN | 38 | 0.3 | 37.7 | 5.709 | 4.225 | 23.3 | 1.39E-06 | 4.11E-05 | 4.38669 |
| RL27_HUMAN | 30.7 | 0 | 30.7 | Inf | 7.197 | 22.9 | 1.71E-06 | 4.93E-05 | 4.30742 |
| SRP72_HUMAN | 27.7 | 0 | 27.7 | Inf | 7.04 | 22.24 | 2.41E-06 | 6.51E-05 | 4.18635 |
| RLA2_HUMAN | 36 | 0.3 | 35.7 | 5.643 | 4.154 | 21.95 | 2.81E-06 | 7.44E-05 | 4.12872 |
| PABP4_HUMAN | 42 | 0.7 | 41.3 | 4.858 | 3.453 | 21.83 | 2.97E-06 | 7.72E-05 | 4.11255 |
| RS9_HUMAN | 28.7 | 0 | 28.7 | Inf | 7.105 | 21.38 | 3.78E-06 | 9.11E-05 | 4.04072 |
| RACK1_HUMAN | 26.3 | 0 | 26.3 | Inf | 6.975 | 21.46 | 3.62E-06 | 9.11E-05 | 4.04072 |
| RL18_HUMAN | 75.3 | 2.3 | 73 | 3.002 | 2.534 | 21.38 | 3.78E-06 | 9.11E-05 | 4.04072 |
| ILF3_HUMAN | 27 | 0 | 27 | Inf | 7.013 | 21.28 | 3.97E-06 | 9.34E-05 | 4.0297 |
| HNRL1_HUMAN | 25 | 0 | 25 | Inf | 6.898 | 20.55 | 5.81E-06 | 0.0001319 | 3.87976 |
| RRP1B_HUMAN | 26.7 | 0 | 26.7 | Inf | 6.996 | 20.47 | 6.05E-06 | 0.0001326 | 3.87746 |
| HNRPU_HUMAN | 150 | 4.7 | 145.3 | 1.274 | 2.315 | 20.5 | 5.96E-06 | 0.0001326 | 3.87746 |
| RRP5_HUMAN | 27 | 0 | 27 | Inf | 7.014 | 20.42 | 6.21E-06 | 0.0001338 | 3.87354 |
| LYAR_HUMAN | 26.7 | 0 | 26.7 | Inf | 7.002 | 20.02 | 7.66E-06 | 0.0001625 | 3.78915 |
| PRKDC_HUMAN | 50.3 | 1.3 | 49 | 4.112 | 2.758 | 19.96 | 7.90E-06 | 0.0001629 | 3.78808 |
| BCLF1_HUMAN | 23.3 | 0 | 23.3 | Inf | 6.809 | 18.82 | 1.43E-05 | 0.0002848 | 3.54546 |
| HP1B3_HUMAN | 26.7 | 0 | 26.7 | Inf | 6.98 | 18.75 | 1.49E-05 | 0.0002864 | 3.54303 |
| TRIPC_HUMAN | 22.7 | 0 | 22.7 | Inf | 6.751 | 18.73 | 1.51E-05 | 0.0002864 | 3.54303 |
| TOP2A_HUMAN | 22.3 | 0 | 22.3 | Inf | 6.733 | 18.74 | 1.50E-05 | 0.0002864 | 3.54303 |
| HSP7C_HUMAN | 58.7 | 2 | 56.7 | 3.76 | 2.429 | 18.61 | 1.60E-05 | 0.0002993 | 3.52389 |
| POP1_HUMAN | 23 | 0 | 23 | Inf | 6.78 | 18.56 | 1.65E-05 | 0.0003043 | 3.5167 |
| DHX30_HUMAN | 28 | 0 | 28 | Inf | 7.073 | 18.1 | 2.09E-05 | 0.0003745 | 3.42655 |
| RL7_HUMAN | 69 | 2.3 | 66.7 | 1.95 | 2.337 | 18.12 | 2.07E-05 | 0.0003745 | 3.42655 |
| RL7A_HUMAN | 84.3 | 3.7 | 80.6 | 2.026 | 2.015 | 17.44 | 2.97E-05 | 0.0005103 | 3.29217 |
| FAS_HUMAN | 22.3 | 0 | 22.3 | Inf | 6.723 | 17.19 | 3.39E-05 | 0.0005668 | 3.24657 |
| ADT2_HUMAN | 28.7 | 0.3 | 28.4 | 5.305 | 3.823 | 17.19 | 3.39E-05 | 0.0005668 | 3.24657 |
| U2AF5_HUMAN | 20.3 | 0 | 20.3 | Inf | 6.606 | 17.06 | 3.62E-05 | 0.0005975 | 3.22366 |
| PFKAP_HUMAN | 20.3 | 0 | 20.3 | Inf | 6.597 | 16.79 | 4.18E-05 | 0.0006818 | 3.16634 |
| RL26_HUMAN | 86 | 3.7 | 82.3 | 2.825 | 2.131 | 16.53 | 4.80E-05 | 0.0007721 | 3.11233 |
| RRBP1_HUMAN | 22.3 | 0 | 22.3 | Inf | 6.739 | 16.42 | 5.08E-05 | 0.0008077 | 3.09275 |
| HS90B_HUMAN | 131.3 | 6.7 | 124.6 | 2.499 | 1.884 | 16.37 | 5.21E-05 | 0.0008186 | 3.08693 |
| RL5_HUMAN | 21.7 | 0 | 21.7 | Inf | 6.707 | 16.26 | 5.52E-05 | 0.0008489 | 3.07114 |
| ADT3_HUMAN | 19.3 | 0 | 19.3 | Inf | 6.533 | 16.25 | 5.54E-05 | 0.0008489 | 3.07114 |
| TBB4A_HUMAN | 114.3 | 7.3 | 107 | 1.792 | 1.49 | 16.18 | 5.76E-05 | 0.0008718 | 3.05958 |
| RS13_HUMAN | 28 | 0.3 | 27.7 | 5.287 | 3.8 | 16.15 | 5.85E-05 | 0.000875 | 3.05799 |
| TBB5_HUMAN | 173.3 | 14 | 159.3 | 1.51 | 1.214 | 16.11 | 5.99E-05 | 0.0008852 | 3.05296 |
| MATR3_HUMAN | 21.3 | 0 | 21.3 | Inf | 6.655 | 15.8 | 7.05E-05 | 0.001019 | 2.99183 |
| SRSF3_HUMAN | 19.7 | 0 | 19.7 | Inf | 6.566 | 15.8 | 7.03E-05 | 0.001019 | 2.99183 |
| FXR1_HUMAN | 20 | 0 | 20 | Inf | 6.585 | 15.64 | 7.65E-05 | 0.001081 | 2.96617 |
| U2AF2_HUMAN | 19.3 | 0 | 19.3 | Inf | 6.536 | 15.62 | 7.74E-05 | 0.001081 | 2.96617 |
| RS3A_HUMAN | 90.3 | 4 | 86.3 | 2.4 | 2.035 | 15.64 | 7.67E-05 | 0.001081 | 2.96617 |
| NOP58_HUMAN | 18.7 | 0 | 18.7 | Inf | 6.48 | 15.59 | 7.85E-05 | 0.001085 | 2.96457 |
| IF2B3_HUMAN | 19 | 0 | 19 | Inf | 6.515 | 15.56 | 8.00E-05 | 0.001094 | 2.96098 |
| GNL3_HUMAN | 22 | 0 | 22 | Inf | 6.693 | 15.5 | 8.27E-05 | 0.001119 | 2.95117 |
| ATPA_HUMAN | 26 | 0.3 | 25.7 | 5.164 | 3.683 | 15.32 | 9.09E-05 | 0.00121 | 2.91721 |
| TBB4B_HUMAN | 168 | 14 | 154 | 1.818 | 1.169 | 15.25 | 9.43E-05 | 0.001236 | 2.90798 |
| TBB2A_HUMAN | 141.3 | 11 | 130.3 | 1.378 | 1.241 | 15.22 | 9.55E-05 | 0.001239 | 2.90693 |
| NUCL_HUMAN | 149.7 | 9.3 | 140.4 | 1.576 | 1.47 | 15.07 | 0.0001036 | 0.001322 | 2.87877 |
| COPA_HUMAN | 17.7 | 0 | 17.7 | Inf | 6.407 | 15 | 0.0001076 | 0.001356 | 2.86774 |
| SRPK1_HUMAN | 18 | 0 | 18 | Inf | 6.435 | 14.94 | 0.000111 | 0.00137 | 2.86328 |
| RLA0_HUMAN | 43.7 | 1.3 | 42.4 | 3.93 | 2.581 | 14.95 | 0.0001107 | 0.00137 | 2.86328 |
| ATPB_HUMAN | 25.3 | 0.3 | 25 | 5.125 | 3.645 | 14.89 | 0.000114 | 0.001394 | 2.85574 |
| RS24_HUMAN | 25.3 | 0.3 | 25 | 5.13 | 3.649 | 14.51 | 0.0001391 | 0.001685 | 2.7734 |
| LC7L2_HUMAN | 17.7 | 0 | 17.7 | Inf | 6.401 | 14.43 | 0.0001458 | 0.001701 | 2.7693 |
| RL23_HUMAN | 31.7 | 0.7 | 31 | 4.469 | 3.065 | 14.24 | 0.0001611 | 0.001863 | 2.72979 |
| LPPRC_HUMAN | 16.7 | 0 | 16.7 | Inf | 6.318 | 14 | 0.0001828 | 0.002039 | 2.69058 |
| FBRL_HUMAN | 24.7 | 0.3 | 24.4 | 2.289 | 3.565 | 13.88 | 0.0001944 | 0.002132 | 2.67121 |
| RRP12_HUMAN | 17.7 | 0 | 17.7 | Inf | 6.386 | 13.8 | 0.0002031 | 0.002208 | 2.656 |
| AT2A2_HUMAN | 16 | 0 | 16 | Inf | 6.26 | 13.67 | 0.0002179 | 0.00231 | 2.63639 |
| SYEP_HUMAN | 16.3 | 0 | 16.3 | Inf | 6.281 | 13.58 | 0.0002282 | 0.002399 | 2.61997 |
| NOC3L_HUMAN | 16 | 0 | 16 | Inf | 6.255 | 13.31 | 0.0002636 | 0.002748 | 2.56098 |
| RBM28_HUMAN | 15.3 | 0 | 15.3 | Inf | 6.202 | 13.21 | 0.0002779 | 0.002861 | 2.54348 |
| RBM25_HUMAN | 15.3 | 0 | 15.3 | Inf | 6.197 | 13.21 | 0.0002789 | 0.002861 | 2.54348 |
| MBB1A_HUMAN | 17 | 0 | 17 | Inf | 6.34 | 13.09 | 0.0002976 | 0.003004 | 2.5223 |
| CCD86_HUMAN | 15.3 | 0 | 15.3 | Inf | 6.194 | 13.05 | 0.0003036 | 0.003041 | 2.51698 |
| RL31_HUMAN | 35.3 | 1 | 34.3 | 2.436 | 2.605 | 13.01 | 0.0003094 | 0.003074 | 2.5123 |
| RL12_HUMAN | 50.3 | 2 | 48.3 | 1.964 | 2.129 | 12.97 | 0.0003171 | 0.003127 | 2.50487 |
| RL27A_HUMAN | 22.7 | 0.3 | 22.4 | 4.98 | 3.495 | 12.88 | 0.0003319 | 0.003223 | 2.49174 |
| PUF60_HUMAN | 15 | 0 | 15 | Inf | 6.167 | 12.75 | 0.0003553 | 0.003398 | 2.46878 |
| RS4X_HUMAN | 65.3 | 3 | 62.3 | 1.76 | 1.907 | 12.57 | 0.0003924 | 0.003725 | 2.42887 |
| RT22_HUMAN | 15.3 | 0 | 15.3 | Inf | 6.201 | 12.52 | 0.0004022 | 0.003778 | 2.42274 |
| SRSF5_HUMAN | 14.3 | 0 | 14.3 | Inf | 6.102 | 12.51 | 0.000404 | 0.003778 | 2.42274 |
| NOP16_HUMAN | 14.3 | 0 | 14.3 | Inf | 6.107 | 12.45 | 0.0004185 | 0.003886 | 2.4105 |
| SMCA5_HUMAN | 15.3 | 0 | 15.3 | Inf | 6.184 | 12.4 | 0.0004288 | 0.003898 | 2.40916 |
| RL32_HUMAN | 22 | 0.3 | 21.7 | 4.938 | 3.453 | 12.4 | 0.0004284 | 0.003898 | 2.40916 |
| RL10_HUMAN | 56.7 | 3 | 53.7 | 1.909 | 1.751 | 12.4 | 0.000429 | 0.003898 | 2.40916 |
| IF4A1_HUMAN | 26.7 | 0.7 | 26 | 4.205 | 2.804 | 12.22 | 0.0004731 | 0.004238 | 2.37284 |
| RS11_HUMAN | 48.7 | 2.3 | 46.4 | 2.384 | 1.912 | 12.23 | 0.0004702 | 0.004238 | 2.37284 |
| HNRPF_HUMAN | 14 | 0 | 14 | Inf | 6.072 | 12.17 | 0.0004849 | 0.004313 | 2.36522 |
| DYHC1_HUMAN | 14.7 | 0 | 14.7 | Inf | 6.133 | 12.13 | 0.0004949 | 0.00437 | 2.35952 |
| RL30_HUMAN | 38 | 1.3 | 36.7 | 2.415 | 2.332 | 12.12 | 0.0004982 | 0.00437 | 2.35952 |
| EFTU_HUMAN | 14.3 | 0 | 14.3 | Inf | 6.109 | 12.09 | 0.0005068 | 0.004375 | 2.35902 |
| RL3_HUMAN | 107.3 | 6 | 101.3 | 1.646 | 1.61 | 12.08 | 0.000509 | 0.004375 | 2.35902 |
| TR150_HUMAN | 15 | 0 | 15 | Inf | 6.189 | 11.9 | 0.0005607 | 0.004786 | 2.32003 |
| EF1A1_HUMAN | 46.3 | 2.3 | 44 | 3.194 | 1.869 | 11.78 | 0.0005986 | 0.005043 | 2.29731 |
| TBA1B_HUMAN | 132.3 | 11 | 121.3 | 1.543 | 1.131 | 11.73 | 0.0006157 | 0.005152 | 2.28802 |
| SRP68_HUMAN | 13.7 | 0 | 13.7 | Inf | 6.034 | 11.7 | 0.0006238 | 0.005187 | 2.28508 |
| DHX15_HUMAN | 14.7 | 0 | 14.7 | Inf | 6.15 | 11.65 | 0.0006414 | 0.005298 | 2.27589 |
| PGAM5_HUMAN | 14.7 | 0 | 14.7 | Inf | 6.146 | 11.59 | 0.0006632 | 0.005407 | 2.26704 |
| LARP1_HUMAN | 14.3 | 0 | 14.3 | Inf | 6.118 | 11.59 | 0.0006631 | 0.005407 | 2.26704 |
| RBM39_HUMAN | 13.3 | 0 | 13.3 | Inf | 6.006 | 11.54 | 0.0006804 | 0.005513 | 2.25861 |
| XRCC5_HUMAN | 20.3 | 0.3 | 20 | 4.804 | 3.33 | 11.38 | 0.0007431 | 0.005982 | 2.22315 |
| NUMA1_HUMAN | 13.3 | 0 | 13.3 | Inf | 5.998 | 11.3 | 0.0007738 | 0.00619 | 2.20831 |
| HNRPQ_HUMAN | 14.7 | 0 | 14.7 | Inf | 6.162 | 11.2 | 0.0008198 | 0.006518 | 2.18589 |
| MCM7_HUMAN | 13 | 0 | 13 | Inf | 5.96 | 11.17 | 0.0008333 | 0.006584 | 2.18151 |
| XRCC6_HUMAN | 20 | 0.3 | 19.7 | 4.792 | 3.312 | 11.15 | 0.0008392 | 0.006589 | 2.18118 |
| RT09_HUMAN | 13 | 0 | 13 | Inf | 5.966 | 11.05 | 0.0008889 | 0.006937 | 2.15883 |
| SRSF2_HUMAN | 20.7 | 0.3 | 20.4 | 4.832 | 3.359 | 10.89 | 0.0009654 | 0.007442 | 2.12831 |
| STAU1_HUMAN | 12.3 | 0 | 12.3 | Inf | 5.894 | 10.84 | 0.0009949 | 0.007623 | 2.11787 |
| TBA4A_HUMAN | 107 | 8.7 | 98.3 | 1.571 | 1.166 | 10.61 | 0.001122 | 0.008548 | 2.06814 |
| TBA1A_HUMAN | 129 | 11 | 118 | 1.507 | 1.095 | 10.52 | 0.001179 | 0.008875 | 2.05183 |
| PR40A_HUMAN | 12 | 0 | 12 | Inf | 5.853 | 10.36 | 0.001286 | 0.009512 | 2.02173 |
| TBB8_HUMAN | 62.3 | 4 | 58.3 | 1.853 | 1.492 | 10.31 | 0.001326 | 0.009751 | 2.01095 |
| RL38_HUMAN | 12 | 0 | 12 | Inf | 5.855 | 10.26 | 0.001356 | 0.009912 | 2.00384 |
| TBA1C_HUMAN | 116.3 | 9.3 | 107 | 1.475 | 1.162 | 10.25 | 0.001365 | 0.009923 | 2.00336 |
| CAVN1_HUMAN | 11.7 | 0 | 11.7 | Inf | 5.811 | 10.08 | 0.001496 | 0.01063 | 1.97347 |
| NSUN2_HUMAN | 11.7 | 0 | 11.7 | Inf | 5.815 | 10.01 | 0.001561 | 0.01103 | 1.95742 |
| QCR2_HUMAN | 11.7 | 0 | 11.7 | Inf | 5.805 | 9.985 | 0.001578 | 0.01109 | 1.95507 |
| IF2B2_HUMAN | 11.3 | 0 | 11.3 | Inf | 5.775 | 9.9 | 0.001653 | 0.01155 | 1.93742 |
| RRS1_HUMAN | 11.3 | 0 | 11.3 | Inf | 5.779 | 9.808 | 0.001737 | 0.01201 | 1.92046 |
| UHRF1_HUMAN | 11 | 0 | 11 | Inf | 5.732 | 9.785 | 0.001759 | 0.0121 | 1.91721 |
| DDX18_HUMAN | 11 | 0 | 11 | Inf | 5.726 | 9.768 | 0.001776 | 0.01214 | 1.91578 |
| RL9_HUMAN | 12.3 | 0 | 12.3 | Inf | 5.896 | 9.722 | 0.001821 | 0.01239 | 1.90693 |
| RS25_HUMAN | 11.7 | 0 | 11.7 | Inf | 5.819 | 9.623 | 0.001922 | 0.01283 | 1.89177 |
| ERH_HUMAN | 11.3 | 0 | 11.3 | Inf | 5.768 | 9.609 | 0.001936 | 0.01283 | 1.89177 |
| RL36A_HUMAN | 18.7 | 0.3 | 18.4 | 1.906 | 3.167 | 9.632 | 0.001912 | 0.01283 | 1.89177 |
| RS18_HUMAN | 28.3 | 1 | 27.3 | 3.727 | 2.36 | 9.609 | 0.001936 | 0.01283 | 1.89177 |
| DDX3X_HUMAN | 49.3 | 2.7 | 46.6 | 1.79 | 1.713 | 9.618 | 0.001926 | 0.01283 | 1.89177 |
| BMS1_HUMAN | 11.7 | 0 | 11.7 | Inf | 5.814 | 9.574 | 0.001974 | 0.01301 | 1.88572 |
| PRP8_HUMAN | 11 | 0 | 11 | Inf | 5.723 | 9.38 | 0.002194 | 0.01438 | 1.84224 |
| DDX10_HUMAN | 10.7 | 0 | 10.7 | Inf | 5.683 | 9.365 | 0.002212 | 0.01443 | 1.84073 |
| NOM1_HUMAN | 10.3 | 0 | 10.3 | Inf | 5.638 | 9.222 | 0.002391 | 0.01544 | 1.81135 |
| AT1A1_HUMAN | 11.7 | 0 | 11.7 | Inf | 5.788 | 9.14 | 0.0025 | 0.0159 | 1.7986 |
| NOG1_HUMAN | 10 | 0 | 10 | Inf | 5.599 | 8.896 | 0.002857 | 0.0179 | 1.74715 |
| DNJC9_HUMAN | 10.3 | 0 | 10.3 | Inf | 5.63 | 8.843 | 0.002942 | 0.01835 | 1.73636 |
| EF1G_HUMAN | 10 | 0 | 10 | Inf | 5.589 | 8.823 | 0.002974 | 0.01845 | 1.734 |
| SPS2L_HUMAN | 10.3 | 0 | 10.3 | Inf | 5.634 | 8.783 | 0.00304 | 0.01877 | 1.72654 |
| KRR1_HUMAN | 11 | 0 | 11 | Inf | 5.708 | 8.755 | 0.003087 | 0.01897 | 1.72193 |
| NOG2_HUMAN | 9.7 | 0 | 9.7 | Inf | 5.545 | 8.674 | 0.003229 | 0.01974 | 1.70465 |
| TCPD_HUMAN | 9.7 | 0 | 9.7 | Inf | 5.547 | 8.556 | 0.003443 | 0.02096 | 1.67861 |
| SSF1_HUMAN | 9.3 | 0 | 9.3 | Inf | 5.496 | 8.413 | 0.003726 | 0.02257 | 1.64647 |
| RENT1_HUMAN | 9.3 | 0 | 9.3 | Inf | 5.501 | 8.378 | 0.003798 | 0.0229 | 1.64016 |
| EBP2_HUMAN | 11 | 0 | 11 | Inf | 5.76 | 8.227 | 0.004126 | 0.02476 | 1.60625 |
| PARP1_HUMAN | 11 | 0 | 11 | Inf | 5.755 | 8.182 | 0.004231 | 0.02526 | 1.59757 |
| RL35A_HUMAN | 15.7 | 0.3 | 15.4 | 4.439 | 2.964 | 8.172 | 0.004255 | 0.02529 | 1.59705 |
| FXR2_HUMAN | 9.3 | 0 | 9.3 | Inf | 5.491 | 8.122 | 0.004373 | 0.02563 | 1.59125 |
| MOV10_HUMAN | 9 | 0 | 9 | Inf | 5.446 | 8.128 | 0.004359 | 0.02563 | 1.59125 |
| ROA0_HUMAN | 15.7 | 0.3 | 15.4 | 4.446 | 2.968 | 8.136 | 0.004338 | 0.02563 | 1.59125 |
| RL22L_HUMAN | 9 | 0 | 9 | Inf | 5.446 | 7.973 | 0.004749 | 0.02758 | 1.55941 |
| TCPG_HUMAN | 9 | 0 | 9 | Inf | 5.437 | 7.881 | 0.004995 | 0.02862 | 1.54333 |
| DDX54_HUMAN | 16 | 0.3 | 15.7 | 1.685 | 2.946 | 7.887 | 0.004978 | 0.02862 | 1.54333 |
| QCR1_HUMAN | 9 | 0 | 9 | Inf | 5.436 | 7.869 | 0.005029 | 0.02869 | 1.54227 |
| U520_HUMAN | 10 | 0 | 10 | Inf | 5.596 | 7.822 | 0.00516 | 0.0293 | 1.53313 |
| RS7_HUMAN | 34.3 | 1.7 | 32.6 | 3.266 | 1.952 | 7.792 | 0.005248 | 0.02967 | 1.52768 |
| DDX47_HUMAN | 8.7 | 0 | 8.7 | Inf | 5.397 | 7.746 | 0.005384 | 0.0303 | 1.51856 |
| RBMX_HUMAN | 20 | 0.7 | 19.3 | 3.794 | 2.4 | 7.732 | 0.005424 | 0.03039 | 1.51727 |
| TCPA_HUMAN | 8.7 | 0 | 8.7 | Inf | 5.397 | 7.699 | 0.005526 | 0.03083 | 1.51103 |
| RS27_HUMAN | 20.3 | 0.7 | 19.6 | 1.808 | 2.365 | 7.677 | 0.005593 | 0.03107 | 1.50766 |
| SDA1_HUMAN | 8.7 | 0 | 8.7 | Inf | 5.389 | 7.636 | 0.00572 | 0.03164 | 1.49976 |
| SMN_HUMAN | 8.3 | 0 | 8.3 | Inf | 5.333 | 7.556 | 0.005981 | 0.03277 | 1.48452 |
| HNRPR_HUMAN | 15 | 0.3 | 14.7 | 4.393 | 2.912 | 7.55 | 0.006002 | 0.03277 | 1.48452 |
| H90B2_HUMAN | 23.7 | 1 | 22.7 | 3.435 | 2.078 | 7.538 | 0.00604 | 0.03283 | 1.48373 |
| SYRC_HUMAN | 8.3 | 0 | 8.3 | Inf | 5.338 | 7.508 | 0.006143 | 0.03325 | 1.47821 |
| NHP2_HUMAN | 8.3 | 0 | 8.3 | Inf | 5.344 | 7.489 | 0.006209 | 0.03332 | 1.4773 |
| MK67I_HUMAN | 8.3 | 0 | 8.3 | Inf | 5.331 | 7.489 | 0.006208 | 0.03332 | 1.4773 |
| BRX1_HUMAN | 8.3 | 0 | 8.3 | Inf | 5.338 | 7.441 | 0.006376 | 0.03408 | 1.4675 |
| RL26L_HUMAN | 49.7 | 3 | 46.7 | 2.219 | 1.621 | 7.423 | 0.006438 | 0.03424 | 1.46547 |
| TBB6_HUMAN | 38.3 | 2.3 | 36 | 2.04 | 1.571 | 7.417 | 0.006461 | 0.03424 | 1.46547 |
| LUC7L_HUMAN | 8.3 | 0 | 8.3 | Inf | 5.336 | 7.407 | 0.006498 | 0.0343 | 1.46471 |
| SPB1_HUMAN | 8.3 | 0 | 8.3 | Inf | 5.349 | 7.357 | 0.006679 | 0.03511 | 1.45457 |
| IF4G1_HUMAN | 8.3 | 0 | 8.3 | Inf | 5.324 | 7.335 | 0.006764 | 0.03541 | 1.45087 |
| TSR1_HUMAN | 8.3 | 0 | 8.3 | Inf | 5.353 | 7.304 | 0.00688 | 0.03587 | 1.44527 |
| DDX52_HUMAN | 8 | 0 | 8 | Inf | 5.287 | 7.159 | 0.007457 | 0.03773 | 1.42331 |
| DDX56_HUMAN | 8 | 0 | 8 | Inf | 5.274 | 7.167 | 0.007424 | 0.03773 | 1.42331 |
| AP3D1_HUMAN | 9.3 | 0 | 9.3 | Inf | 5.489 | 7.136 | 0.007556 | 0.03799 | 1.42033 |
| PCBP2_HUMAN | 8 | 0 | 8 | Inf | 5.281 | 7.121 | 0.007619 | 0.03816 | 1.41839 |
| TF3C1_HUMAN | 8 | 0 | 8 | Inf | 5.279 | 7.075 | 0.007816 | 0.03884 | 1.41072 |
| CC137_HUMAN | 8 | 0 | 8 | Inf | 5.271 | 7.068 | 0.007847 | 0.03884 | 1.41072 |
| SYMC_HUMAN | 8 | 0 | 8 | Inf | 5.293 | 6.957 | 0.008351 | 0.04058 | 1.39169 |
| RL22_HUMAN | 8 | 0 | 8 | Inf | 5.282 | 6.955 | 0.008359 | 0.04058 | 1.39169 |
| DKC1_HUMAN | 7.7 | 0 | 7.7 | Inf | 5.223 | 6.957 | 0.008348 | 0.04058 | 1.39169 |
| RS20_HUMAN | 14.3 | 0.3 | 14 | 4.29 | 2.831 | 6.885 | 0.008691 | 0.04203 | 1.37644 |
| RSSA_HUMAN | 7.7 | 0 | 7.7 | Inf | 5.218 | 6.814 | 0.009043 | 0.0434 | 1.36251 |
| TF3C3_HUMAN | 7.7 | 0 | 7.7 | Inf | 5.213 | 6.777 | 0.009235 | 0.044 | 1.35655 |
| SNUT1_HUMAN | 7.7 | 0 | 7.7 | Inf | 5.209 | 6.78 | 0.009217 | 0.044 | 1.35655 |
| RS23_HUMAN | 34.7 | 2 | 32.7 | 1.424 | 1.591 | 6.702 | 0.009631 | 0.04571 | 1.33999 |
| SLTM_HUMAN | 7.3 | 0 | 7.3 | Inf | 5.167 | 6.538 | 0.01056 | 0.04961 | 1.30443 |
| TBA4B_HUMAN | 7.3 | 0 | 7.3 | Inf | 5.162 | 6.536 | 0.01057 | 0.04961 | 1.30443 |

**Table S2: List of significantly enriched proteins in SINV-DDX5_IP versus SINV-IgG_IP**

**1st column** : Protein name

**2nd column** : Column named as the treatment level with the mean raw spectral counts observed for this condition

**3rd column** : Column named as the control level with the mean raw spectral counts observed for this condition

**deltaSC** : Difference between mean raw spectral counts

**lFC.Av** : Log fold change computed from the mean expression levels taking into account the given normalization factors.

**logFC** : Log fold change estimated by fitting the given GLM model. The reference level of the main factor is taken as control.

**LR** : The likelihood ratio for edgeR.

**p.val** : The unadjusted p-values obtained from the tests.

**adjp** : The multitest adjusted p-values with FDR control.

|  | SINV_DDX5IP | SINV_IgGIP | deltaSC | lFC_Av | LogFC | LR | p_value | adjp | negLog_adjp |
| --- | --- | --- | --- | --- | --- | --- | --- | --- | --- |
| DDX5_HUMAN | 717.3 | 0.3 | 717 | 9.588 | 8.422 | 157.9 | 3.17E-36 | 3.87E-33 | 32.41274 |
| DDX21_HUMAN | 137.3 | 0.7 | 136.6 | 6.197 | 5.12 | 55.9 | 7.61E-14 | 4.64E-11 | 10.33386 |
| DDX17_HUMAN | 115.3 | 0.3 | 115 | 6.94 | 5.776 | 54.24 | 1.77E-13 | 7.20E-11 | 10.14285 |
| RL6_HUMAN | 89.7 | 0.3 | 89.4 | 6.578 | 5.414 | 44.59 | 2.42E-11 | 7.38E-09 | 8.13183 |
| YBOX1_HUMAN | 138 | 2 | 136 | 3.513 | 3.568 | 40.57 | 1.90E-10 | 4.62E-08 | 7.33564 |
| SRRM2_HUMAN | 66.3 | 0 | 66.3 | Inf | 8.314 | 37.75 | 8.05E-10 | 1.64E-07 | 6.78648 |
| YBOX3_HUMAN | 51 | 0 | 51 | Inf | 7.924 | 32.25 | 1.36E-08 | 1.84E-06 | 5.73565 |
| RL36_HUMAN | 48.7 | 0 | 48.7 | Inf | 7.849 | 31.72 | 1.78E-08 | 2.08E-06 | 5.68298 |
| SRRM1_HUMAN | 42.7 | 0 | 42.7 | Inf | 7.665 | 28.78 | 8.11E-08 | 7.60E-06 | 5.11942 |
| KI67_HUMAN | 47.3 | 0 | 47.3 | Inf | 7.816 | 28.19 | 1.10E-07 | 8.83E-06 | 5.05384 |
| RL35_HUMAN | 43 | 0 | 43 | Inf | 7.684 | 28.2 | 1.09E-07 | 8.83E-06 | 5.05384 |
| RS6_HUMAN | 84 | 1.7 | 82.3 | 4.158 | 3.167 | 28.09 | 1.16E-07 | 8.83E-06 | 5.05384 |
| RL21_HUMAN | 42.7 | 0 | 42.7 | Inf | 7.669 | 27.36 | 1.69E-07 | 1.14E-05 | 4.94271 |
| NOP56_HUMAN | 54 | 0.3 | 53.7 | 5.883 | 4.728 | 24.71 | 6.66E-07 | 3.90E-05 | 4.40905 |
| RL18A_HUMAN | 62.7 | 1 | 61.7 | 4.458 | 3.428 | 24.69 | 6.72E-07 | 3.90E-05 | 4.40905 |
| PNPT1_HUMAN | 48 | 0 | 48 | Inf | 7.879 | 23.35 | 1.35E-06 | 6.84E-05 | 4.16494 |
| RS13_HUMAN | 29.7 | 0 | 29.7 | Inf | 7.131 | 21.28 | 3.96E-06 | 0.0001776 | 3.75056 |
| RL13A_HUMAN | 47.7 | 0.7 | 47 | 4.675 | 3.604 | 21.23 | 4.07E-06 | 0.0001776 | 3.75056 |
| RL17_HUMAN | 56.7 | 1 | 55.7 | 4.349 | 3.321 | 21.31 | 3.91E-06 | 0.0001776 | 3.75056 |
| PABP1_HUMAN | 70.7 | 1.7 | 69 | 3.932 | 2.958 | 21.16 | 4.23E-06 | 0.0001776 | 3.75056 |
| Pol_S_SINV_Capside | 119 | 5 | 114 | 2.229 | 2.061 | 21.19 | 4.16E-06 | 0.0001776 | 3.75056 |
| NUCL_HUMAN | 163.3 | 5.3 | 158 | 3.446 | 2.715 | 20.72 | 5.31E-06 | 0.0002086 | 3.68069 |
| LYAR_HUMAN | 28.3 | 0 | 28.3 | Inf | 7.069 | 20.37 | 6.38E-06 | 0.0002429 | 3.61457 |
| RS4X_HUMAN | 72.3 | 2 | 70.3 | 3.696 | 2.739 | 19.93 | 8.02E-06 | 0.0002958 | 3.529 |
| NAT10_HUMAN | 36.3 | 0.3 | 36 | 5.275 | 4.117 | 19.51 | 9.99E-06 | 0.0003405 | 3.46788 |
| NOP2_HUMAN | 53.3 | 1 | 52.3 | 4.262 | 3.238 | 19.5 | 1.01E-05 | 0.0003405 | 3.46788 |
| POP1_HUMAN | 26.3 | 0 | 26.3 | Inf | 6.975 | 19.05 | 1.28E-05 | 0.00042 | 3.37675 |
| RL5_HUMAN | 26.7 | 0 | 26.7 | Inf | 6.983 | 18.62 | 1.60E-05 | 0.0004856 | 3.31372 |
| BCLF1_HUMAN | 25 | 0 | 25 | Inf | 6.886 | 18.67 | 1.55E-05 | 0.0004856 | 3.31372 |
| TOP1_HUMAN | 35 | 0.3 | 34.7 | 5.226 | 4.068 | 18.64 | 1.58E-05 | 0.0004856 | 3.31372 |
| RRP1B_HUMAN | 26.3 | 0 | 26.3 | Inf | 6.961 | 18.56 | 1.65E-05 | 0.0004885 | 3.31114 |
| SRP72_HUMAN | 23.7 | 0 | 23.7 | Inf | 6.818 | 17.76 | 2.51E-05 | 0.0006977 | 3.15633 |
| HSP7C_HUMAN | 67.3 | 2 | 65.3 | 3.108 | 2.597 | 17.75 | 2.52E-05 | 0.0006977 | 3.15633 |
| HNRPU_HUMAN | 166 | 9 | 157 | 1.96 | 1.782 | 17.83 | 2.41E-05 | 0.0006977 | 3.15633 |
| RRP12_HUMAN | 22.7 | 0 | 22.7 | Inf | 6.75 | 17.31 | 3.18E-05 | 0.0008066 | 3.09334 |
| RL10_HUMAN | 63.7 | 2 | 61.7 | 3.478 | 2.518 | 17.32 | 3.16E-05 | 0.0008066 | 3.09334 |
| MBB1A_HUMAN | 23.7 | 0 | 23.7 | Inf | 6.841 | 16.86 | 4.03E-05 | 0.0009999 | 3.00004 |
| RL10A_HUMAN | 63.7 | 2 | 61.7 | 3.506 | 2.553 | 16.82 | 4.11E-05 | 0.0009999 | 3.00004 |
| DHX30_HUMAN | 23.3 | 0 | 23.3 | Inf | 6.793 | 16.66 | 4.46E-05 | 0.001066 | 2.97224 |
| RS7_HUMAN | 31.3 | 0.3 | 31 | 5.071 | 3.914 | 16.55 | 4.73E-05 | 0.001108 | 2.95546 |
| GNL3_HUMAN | 21.7 | 0 | 21.7 | Inf | 6.696 | 16.37 | 5.22E-05 | 0.001178 | 2.92885 |
| NOP58_HUMAN | 21.3 | 0 | 21.3 | Inf | 6.661 | 16.4 | 5.14E-05 | 0.001178 | 2.92885 |
| ATPB_HUMAN | 21.3 | 0 | 21.3 | Inf | 6.66 | 16.18 | 5.76E-05 | 0.001274 | 2.89483 |
| RRP5_HUMAN | 21 | 0 | 21 | Inf | 6.634 | 16.14 | 5.89E-05 | 0.001281 | 2.89245 |
| RL7A_HUMAN | 90.3 | 3.7 | 86.6 | 3.135 | 2.28 | 16.09 | 6.05E-05 | 0.001293 | 2.8884 |
| NUMA1_HUMAN | 22.7 | 0 | 22.7 | Inf | 6.727 | 15.92 | 6.62E-05 | 0.001319 | 2.87976 |
| RL27_HUMAN | 29.3 | 0.3 | 29 | 4.965 | 3.81 | 15.52 | 8.16E-05 | 0.001553 | 2.80883 |
| RBM28_HUMAN | 21.3 | 0 | 21.3 | Inf | 6.69 | 15.29 | 9.24E-05 | 0.001679 | 2.77495 |
| U2AF5_HUMAN | 19.3 | 0 | 19.3 | Inf | 6.524 | 15.08 | 0.0001033 | 0.00185 | 2.73283 |
| TOP2A_HUMAN | 20 | 0 | 20 | Inf | 6.594 | 15.03 | 0.000106 | 0.001871 | 2.72793 |
| PABP4_HUMAN | 36 | 0.7 | 35.3 | 4.281 | 3.215 | 14.97 | 0.0001091 | 0.001871 | 2.72793 |
| DDX24_HUMAN | 31 | 0.3 | 30.7 | 5.025 | 3.882 | 14.84 | 0.000117 | 0.00198 | 2.70333 |
| RL27A_HUMAN | 28.7 | 0.3 | 28.4 | 4.906 | 3.761 | 14.78 | 0.0001206 | 0.002012 | 2.69637 |
| RL7_HUMAN | 70.3 | 2.3 | 68 | 3.46 | 2.571 | 14.65 | 0.0001294 | 0.002129 | 2.67182 |
| MATR3_HUMAN | 18.7 | 0 | 18.7 | Inf | 6.477 | 14.4 | 0.000148 | 0.002404 | 2.61907 |
| HP1B3_HUMAN | 22 | 0 | 22 | Inf | 6.744 | 14.3 | 0.0001561 | 0.002447 | 2.61137 |
| DHX15_HUMAN | 19.7 | 0 | 19.7 | Inf | 6.559 | 14.31 | 0.000155 | 0.002447 | 2.61137 |
| RLA2_HUMAN | 40.3 | 1 | 39.3 | 2.513 | 2.76 | 14.29 | 0.0001567 | 0.002447 | 2.61137 |
| IF2B3_HUMAN | 18.3 | 0 | 18.3 | Inf | 6.458 | 14.18 | 0.0001662 | 0.002499 | 2.60223 |
| RRBP1_HUMAN | 18 | 0 | 18 | Inf | 6.431 | 14.01 | 0.0001819 | 0.002702 | 2.56831 |
| RL18_HUMAN | 68 | 2.7 | 65.3 | 3.203 | 2.306 | 13.98 | 0.0001846 | 0.002709 | 2.56719 |
| FXR1_HUMAN | 17.7 | 0 | 17.7 | Inf | 6.405 | 13.85 | 0.0001982 | 0.002874 | 2.54151 |
| SRSF3_HUMAN | 17.7 | 0 | 17.7 | Inf | 6.389 | 13.75 | 0.0002087 | 0.002967 | 2.52768 |
| RS11_HUMAN | 48.7 | 1.7 | 47 | 3.385 | 2.407 | 13.69 | 0.0002156 | 0.003018 | 2.52028 |
| EBP2_HUMAN | 17.3 | 0 | 17.3 | Inf | 6.387 | 13.42 | 0.0002493 | 0.0033 | 2.48149 |
| AT2A2_HUMAN | 17 | 0 | 17 | Inf | 6.342 | 13.42 | 0.000249 | 0.0033 | 2.48149 |
| RL4_HUMAN | 188.3 | 7.3 | 181 | 3.203 | 2.664 | 13.42 | 0.0002483 | 0.0033 | 2.48149 |
| RRS1_HUMAN | 17.7 | 0 | 17.7 | Inf | 6.401 | 13.37 | 0.0002561 | 0.003335 | 2.4769 |
| COPA_HUMAN | 17 | 0 | 17 | Inf | 6.351 | 13.36 | 0.0002574 | 0.003335 | 2.4769 |
| RL26_HUMAN | 97 | 5 | 92 | 2.583 | 1.953 | 13.12 | 0.0002916 | 0.003738 | 2.42736 |
| SMCA5_HUMAN | 17.7 | 0 | 17.7 | Inf | 6.418 | 13.09 | 0.0002962 | 0.003758 | 2.42504 |
| RL3_HUMAN | 111.3 | 6.7 | 104.6 | 1.909 | 1.63 | 13.02 | 0.0003086 | 0.003835 | 2.41623 |
| SRP68_HUMAN | 16.7 | 0 | 16.7 | Inf | 6.333 | 12.83 | 0.000341 | 0.004195 | 2.37727 |
| PRKDC_HUMAN | 50.3 | 2 | 48.3 | 3.164 | 2.204 | 12.62 | 0.0003822 | 0.004609 | 2.33639 |
| NOP16_HUMAN | 16 | 0 | 16 | Inf | 6.264 | 12.53 | 0.0004013 | 0.004793 | 2.31939 |
| DDX3X_HUMAN | 42 | 1.3 | 40.7 | 3.504 | 2.516 | 12.32 | 0.0004475 | 0.005278 | 2.27753 |
| XRCC5_HUMAN | 16.7 | 0 | 16.7 | Inf | 6.275 | 12.22 | 0.0004729 | 0.005433 | 2.26496 |
| BMS1_HUMAN | 15.7 | 0 | 15.7 | Inf | 6.243 | 12.23 | 0.0004712 | 0.005433 | 2.26496 |
| RLA0_HUMAN | 44 | 1.7 | 42.3 | 3.227 | 2.246 | 12.1 | 0.0005041 | 0.005739 | 2.24116 |
| HNRPQ_HUMAN | 15.7 | 0 | 15.7 | Inf | 6.214 | 12.05 | 0.0005183 | 0.005845 | 2.23322 |
| RS3A_HUMAN | 94.7 | 5.3 | 89.4 | 2.457 | 1.819 | 12.02 | 0.000525 | 0.005867 | 2.23158 |
| DKC1_HUMAN | 15.3 | 0 | 15.3 | Inf | 6.198 | 11.98 | 0.0005368 | 0.005944 | 2.22592 |
| SRPK1_HUMAN | 15.7 | 0 | 15.7 | Inf | 6.235 | 11.94 | 0.0005507 | 0.006043 | 2.21875 |
| CCD86_HUMAN | 15.3 | 0 | 15.3 | Inf | 6.209 | 11.85 | 0.0005762 | 0.006266 | 2.20301 |
| RL36A_HUMAN | 23 | 0.3 | 22.7 | 4.619 | 3.467 | 11.77 | 0.000602 | 0.006488 | 2.18789 |
| Pol_NS_SINV_nsP2 | 28.7 | 0.7 | 28 | 3.922 | 2.86 | 11.75 | 0.0006083 | 0.006499 | 2.18715 |
| SPS2L_HUMAN | 14.7 | 0 | 14.7 | Inf | 6.141 | 11.69 | 0.0006292 | 0.006664 | 2.17627 |
| RL12_HUMAN | 40 | 1.3 | 38.7 | 3.439 | 2.448 | 11.67 | 0.000635 | 0.006667 | 2.17607 |
| IMA1_HUMAN | 22.7 | 0.3 | 22.4 | 4.597 | 3.445 | 11.64 | 0.0006443 | 0.006707 | 2.17347 |
| PFKAP_HUMAN | 22.7 | 0.3 | 22.4 | 4.603 | 3.449 | 11.61 | 0.0006544 | 0.006755 | 2.17037 |
| TR150_HUMAN | 15.3 | 0 | 15.3 | Inf | 6.192 | 11.56 | 0.0006725 | 0.006883 | 2.16222 |
| RL14_HUMAN | 37.7 | 1 | 36.7 | 1.746 | 2.604 | 11.54 | 0.0006822 | 0.006924 | 2.15964 |
| RL1D1_HUMAN | 46 | 1.7 | 44.3 | 3.292 | 2.337 | 11.51 | 0.0006941 | 0.006987 | 2.15571 |
| U2AF2_HUMAN | 15 | 0 | 15 | Inf | 6.189 | 11.47 | 0.000709 | 0.00702 | 2.15366 |
| NOG1_HUMAN | 14.7 | 0 | 14.7 | Inf | 6.16 | 11.37 | 0.0007483 | 0.007291 | 2.13721 |
| MCM7_HUMAN | 14.7 | 0 | 14.7 | Inf | 6.109 | 11.34 | 0.0007598 | 0.007345 | 2.13401 |
| HNRPR_HUMAN | 14.7 | 0 | 14.7 | Inf | 6.112 | 11.25 | 0.0007953 | 0.007509 | 2.12442 |
| RBMX_HUMAN | 22 | 0.3 | 21.7 | 4.544 | 3.395 | 11.26 | 0.0007916 | 0.007509 | 2.12442 |
| RL26L_HUMAN | 58.7 | 3 | 55.7 | 2.46 | 1.847 | 10.92 | 0.0009535 | 0.008602 | 2.0654 |
| DDX54_HUMAN | 13.3 | 0 | 13.3 | Inf | 6 | 10.83 | 0.0009971 | 0.008868 | 2.05217 |
| RS16_HUMAN | 37.3 | 1.3 | 36 | 3.292 | 2.3 | 10.83 | 0.0009975 | 0.008868 | 2.05217 |
| PUF60_HUMAN | 13.3 | 0 | 13.3 | Inf | 6.003 | 10.8 | 0.001014 | 0.00895 | 2.04818 |
| HS90A_HUMAN | 71.7 | 3.7 | 68 | 2.778 | 1.945 | 10.67 | 0.001087 | 0.009527 | 2.02104 |
| SRSF5_HUMAN | 13.3 | 0 | 13.3 | Inf | 6.02 | 10.56 | 0.001158 | 0.01 | 2 |
| DDX18_HUMAN | 13 | 0 | 13 | Inf | 5.96 | 10.48 | 0.00121 | 0.01031 | 1.98674 |
| EFTU_HUMAN | 13 | 0 | 13 | Inf | 5.947 | 10.47 | 0.00121 | 0.01031 | 1.98674 |
| RT22_HUMAN | 13.3 | 0 | 13.3 | Inf | 6.019 | 10.45 | 0.001226 | 0.01037 | 1.98422 |
| KRR1_HUMAN | 13.3 | 0 | 13.3 | Inf | 5.993 | 10.32 | 0.001313 | 0.01086 | 1.96417 |
| RL30_HUMAN | 37.7 | 1.3 | 36.4 | 3.36 | 2.38 | 10.32 | 0.001319 | 0.01086 | 1.96417 |
| ATD3A_HUMAN | 13.7 | 0 | 13.7 | Inf | 5.992 | 10.15 | 0.001446 | 0.01159 | 1.93592 |
| BRX1_HUMAN | 12.7 | 0 | 12.7 | Inf | 5.908 | 10.07 | 0.001508 | 0.01185 | 1.92628 |
| AT1A1_HUMAN | 12.3 | 0 | 12.3 | Inf | 5.889 | 10.07 | 0.001503 | 0.01185 | 1.92628 |
| RT09_HUMAN | 12.3 | 0 | 12.3 | Inf | 5.891 | 10.04 | 0.001534 | 0.0119 | 1.92445 |
| DNJC9_HUMAN | 12.3 | 0 | 12.3 | Inf | 5.889 | 10.05 | 0.001525 | 0.0119 | 1.92445 |
| CAVN1_HUMAN | 12.3 | 0 | 12.3 | Inf | 5.882 | 10.01 | 0.001554 | 0.01198 | 1.92154 |
| PGAM5_HUMAN | 12.3 | 0 | 12.3 | Inf | 5.881 | 9.983 | 0.00158 | 0.0121 | 1.91721 |
| RL35A_HUMAN | 12.3 | 0 | 12.3 | Inf | 5.902 | 9.905 | 0.001649 | 0.01247 | 1.90413 |
| RACK1_HUMAN | 25.7 | 0.7 | 25 | 3.755 | 2.7 | 9.916 | 0.001638 | 0.01247 | 1.90413 |
| SRSF2_HUMAN | 19.7 | 0.3 | 19.4 | 4.372 | 3.23 | 9.755 | 0.001788 | 0.01344 | 1.8716 |
| DYHC1_HUMAN | 11.7 | 0 | 11.7 | Inf | 5.807 | 9.634 | 0.00191 | 0.01427 | 1.84558 |
| DHX9_HUMAN | 30.3 | 1 | 29.3 | 3.455 | 2.438 | 9.558 | 0.001991 | 0.01479 | 1.83003 |
| RL8_HUMAN | 58 | 3.3 | 54.7 | 1.867 | 1.633 | 9.484 | 0.002072 | 0.0153 | 1.81531 |
| LC7L2_HUMAN | 19.7 | 0.3 | 19.4 | 4.405 | 3.256 | 9.449 | 0.002113 | 0.0155 | 1.80967 |
| IF2B2_HUMAN | 11.7 | 0 | 11.7 | Inf | 5.797 | 9.38 | 0.002194 | 0.016 | 1.79588 |
| TRIPC_HUMAN | 19 | 0.3 | 18.7 | 4.341 | 3.192 | 9.368 | 0.002208 | 0.01601 | 1.79561 |
| STAU1_HUMAN | 11.3 | 0 | 11.3 | Inf | 5.773 | 9.182 | 0.002445 | 0.01762 | 1.75399 |
| NOM1_HUMAN | 11 | 0 | 11 | Inf | 5.726 | 9.124 | 0.002522 | 0.01807 | 1.74304 |
| RL38_HUMAN | 11.3 | 0 | 11.3 | Inf | 5.752 | 9.112 | 0.002539 | 0.01809 | 1.74256 |
| SPB1_HUMAN | 11.7 | 0 | 11.7 | Inf | 5.832 | 9.09 | 0.00257 | 0.01812 | 1.74184 |
| RL23_HUMAN | 29.7 | 1 | 28.7 | 3.362 | 2.355 | 9.087 | 0.002574 | 0.01812 | 1.74184 |
| U520_HUMAN | 11 | 0 | 11 | Inf | 5.742 | 8.984 | 0.002724 | 0.01907 | 1.71965 |
| RL36L_HUMAN | 18.3 | 0.3 | 18 | 4.312 | 3.159 | 8.867 | 0.002904 | 0.02021 | 1.69443 |
| LARP1_HUMAN | 11 | 0 | 11 | Inf | 5.708 | 8.818 | 0.002982 | 0.02064 | 1.68529 |
| SSF1_HUMAN | 10.7 | 0 | 10.7 | Inf | 5.693 | 8.806 | 0.003003 | 0.02067 | 1.68466 |
| DDX10_HUMAN | 10.7 | 0 | 10.7 | Inf | 5.698 | 8.705 | 0.003174 | 0.02172 | 1.66314 |
| SYRC_HUMAN | 10.3 | 0 | 10.3 | Inf | 5.651 | 8.522 | 0.003509 | 0.02374 | 1.62452 |
| RRP1_HUMAN | 10 | 0 | 10 | Inf | 5.596 | 8.366 | 0.003824 | 0.02573 | 1.58956 |
| TSR1_HUMAN | 10 | 0 | 10 | Inf | 5.58 | 8.269 | 0.004034 | 0.02699 | 1.5688 |
| MK67I_HUMAN | 10 | 0 | 10 | Inf | 5.594 | 8.252 | 0.004071 | 0.0271 | 1.56703 |
| HNRL1_HUMAN | 18 | 0.3 | 17.7 | 4.305 | 3.155 | 8.238 | 0.004102 | 0.02715 | 1.56623 |
| SDA1_HUMAN | 10 | 0 | 10 | Inf | 5.595 | 8.123 | 0.004371 | 0.02854 | 1.54455 |
| NHP2_HUMAN | 9.7 | 0 | 9.7 | Inf | 5.545 | 8.119 | 0.004381 | 0.02854 | 1.54455 |
| RS9_HUMAN | 31.3 | 1.3 | 30 | 3.032 | 2.049 | 8.087 | 0.004458 | 0.02888 | 1.5394 |
| RT18B_HUMAN | 9.7 | 0 | 9.7 | Inf | 5.538 | 8.045 | 0.004562 | 0.02935 | 1.53239 |
| RS18_HUMAN | 30.3 | 1.3 | 29 | 3.022 | 2.025 | 8.039 | 0.004578 | 0.02935 | 1.53239 |
| XRCC6_HUMAN | 21.7 | 0.7 | 21 | 3.508 | 2.456 | 7.896 | 0.004954 | 0.03159 | 1.50045 |
| FIP1_HUMAN | 9.7 | 0 | 9.7 | Inf | 5.551 | 7.833 | 0.00513 | 0.03187 | 1.49662 |
| IF4G1_HUMAN | 9.3 | 0 | 9.3 | Inf | 5.501 | 7.806 | 0.005208 | 0.03187 | 1.49662 |
| UHRF1_HUMAN | 9.3 | 0 | 9.3 | Inf | 5.499 | 7.808 | 0.005203 | 0.03187 | 1.49662 |
| TCPG_HUMAN | 9.3 | 0 | 9.3 | Inf | 5.486 | 7.853 | 0.005074 | 0.03187 | 1.49662 |
| ADT2_HUMAN | 30.7 | 1.3 | 29.4 | 3.004 | 2.021 | 7.818 | 0.005172 | 0.03187 | 1.49662 |
| TCPD_HUMAN | 10 | 0 | 10 | Inf | 5.565 | 7.679 | 0.005586 | 0.03368 | 1.47263 |
| SYEP_HUMAN | 16.3 | 0.3 | 16 | 4.091 | 2.959 | 7.602 | 0.005832 | 0.03459 | 1.46105 |
| RS2_HUMAN | 60.7 | 4 | 56.7 | 2.44 | 1.616 | 7.592 | 0.005864 | 0.03459 | 1.46105 |
| NOL10_HUMAN | 9 | 0 | 9 | Inf | 5.453 | 7.523 | 0.006091 | 0.0355 | 1.44977 |
| FBRL_HUMAN | 25 | 1 | 24 | 3.153 | 2.13 | 7.498 | 0.006178 | 0.03583 | 1.44575 |
| NSUN2_HUMAN | 9 | 0 | 9 | Inf | 5.439 | 7.456 | 0.006323 | 0.0365 | 1.43771 |
| TBB4A_HUMAN | 130.7 | 11.3 | 119.4 | 1.217 | 1.087 | 7.417 | 0.00646 | 0.03712 | 1.43039 |
| RPF2_HUMAN | 9 | 0 | 9 | Inf | 5.427 | 7.357 | 0.006679 | 0.0376 | 1.42481 |
| RS23_HUMAN | 33 | 1.7 | 31.3 | 2.838 | 1.875 | 7.223 | 0.007199 | 0.04022 | 1.39556 |
| PR40A_HUMAN | 8.7 | 0 | 8.7 | Inf | 5.367 | 7.196 | 0.007306 | 0.04063 | 1.39115 |
| RS27_HUMAN | 20.3 | 0.7 | 19.6 | 3.446 | 2.389 | 7.17 | 0.007415 | 0.04105 | 1.38669 |
| RL22L_HUMAN | 8.7 | 0 | 8.7 | Inf | 5.371 | 7.135 | 0.00756 | 0.04115 | 1.38563 |
| PRP6_HUMAN | 8.7 | 0 | 8.7 | Inf | 5.367 | 7.136 | 0.007557 | 0.04115 | 1.38563 |
| KRI1_HUMAN | 8.7 | 0 | 8.7 | Inf | 5.39 | 7.093 | 0.007737 | 0.04188 | 1.37799 |
| RS24_HUMAN | 20 | 0.7 | 19.3 | 3.428 | 2.368 | 7.049 | 0.007932 | 0.04275 | 1.36906 |
| CC137_HUMAN | 8.3 | 0 | 8.3 | Inf | 5.35 | 7.015 | 0.008082 | 0.04327 | 1.36381 |
| PESC_HUMAN | 8.3 | 0 | 8.3 | Inf | 5.33 | 7.011 | 0.008101 | 0.04327 | 1.36381 |
| TDIF2_HUMAN | 8.3 | 0 | 8.3 | Inf | 5.337 | 6.953 | 0.008367 | 0.0445 | 1.35164 |
| ILF3_HUMAN | 24.7 | 1 | 23.7 | 3.167 | 2.156 | 6.882 | 0.008709 | 0.04592 | 1.338 |
| LUC7L_HUMAN | 8 | 0 | 8 | Inf | 5.281 | 6.813 | 0.00905 | 0.04732 | 1.32496 |
| F120A_HUMAN | 8 | 0 | 8 | Inf | 5.279 | 6.813 | 0.009052 | 0.04732 | 1.32496 |
| RM11_HUMAN | 8 | 0 | 8 | Inf | 5.283 | 6.784 | 0.009198 | 0.04767 | 1.32175 |
| DDX56_HUMAN | 8 | 0 | 8 | Inf | 5.27 | 6.784 | 0.009198 | 0.04767 | 1.32175 |
| DDX52_HUMAN | 8 | 0 | 8 | Inf | 5.287 | 6.758 | 0.009335 | 0.04818 | 1.31713 |
| RL32_HUMAN | 19.3 | 0.7 | 18.6 | 3.359 | 2.303 | 6.743 | 0.009412 | 0.04837 | 1.31542 |
| BAG2_HUMAN | 8 | 0 | 8 | Inf | 5.298 | 6.672 | 0.009794 | 0.04991 | 1.30181 |

**Table S3: List of significantly enriched proteins in SINV-DDX5_IP versus mock-DDX5_IP**

**1st column** : Protein name

**2nd column** : Column named as the treatment level with the mean raw spectral counts observed for this condition

**3rd column** : Column named as the control level with the mean raw spectral counts observed for this condition

**deltaSC** : Difference between mean raw spectral counts

**lFC.Av** : Log fold change computed from the mean expression levels taking into account the given normalization factors.

**logFC** : Log fold change estimated by fitting the given GLM model. The reference level of the main factor is taken as control.

**LR** : The likelihood ratio for edgeR.

**p.val** : The unadjusted p-values obtained from the tests.

**adjp** : The multitest adjusted p-values with FDR control.

|  | SINV_DDX5IP | mock_DDX5IP | deltaSC | lFC_Av | LogFC | LR | p_value | adjp | negLog_adjp |
| --- | --- | --- | --- | --- | --- | --- | --- | --- | --- |
| Pol_S_SINV_Capside | 119 | 0 | 119 | Inf | 9.879 | 207.9 | 3.90E-47 | 5.36E-44 | 43.27059 |
| Pol_NS_SINV_nsP2 | 28.7 | 0 | 28.7 | Inf | 7.83 | 79.84 | 4.06E-19 | 2.80E-16 | 15.55346 |
| Pol_S_SINV_E1 | 15.3 | 0 | 15.3 | Inf | 6.925 | 47.44 | 5.66E-12 | 2.60E-09 | 8.58503 |
| Pol_S_SINV_E2 | 15 | 0 | 15 | Inf | 6.895 | 46.7 | 8.27E-12 | 2.85E-09 | 8.54561 |
| Pol_NS_SINV_nsP3 | 14.3 | 0 | 14.3 | Inf | 6.862 | 43.42 | 4.42E-11 | 1.22E-08 | 7.91435 |
| MYH9_HUMAN | 349.7 | 0.7 | 349 | 8.624 | 8.477 | 30.21 | 3.87E-08 | 8.88E-06 | 5.05154 |
| MYH10_HUMAN | 84.3 | 0 | 84.3 | Inf | 9.136 | 20.98 | 4.64E-06 | 0.000913 | 3.03953 |
| MYL6_HUMAN | 23.7 | 0.3 | 23.4 | 6.06 | 5.499 | 18.26 | 1.93E-05 | 0.003323 | 2.47847 |
| MYL6B_HUMAN | 5 | 0 | 5 | Inf | 5.292 | 16.9 | 3.93E-05 | 0.006015 | 2.22076 |
| TPM3_HUMAN | 21 | 0 | 21 | Inf | 7.147 | 14.9 | 0.0001132 | 0.01418 | 1.84832 |
| TPM4_HUMAN | 18.7 | 0 | 18.7 | Inf | 6.982 | 14.72 | 0.0001246 | 0.0143 | 1.84466 |
| MYH11_HUMAN | 20.3 | 0.7 | 19.6 | 4.539 | 4.433 | 14.28 | 0.0001573 | 0.01547 | 1.81051 |
| TPM2_HUMAN | 15.3 | 0 | 15.3 | Inf | 6.706 | 14.41 | 0.0001473 | 0.01547 | 1.81051 |
| MYH14_HUMAN | 21 | 0 | 21 | Inf | 7.139 | 13.3 | 0.0002648 | 0.02279 | 1.64226 |
| ML12B_HUMAN | 7.3 | 0 | 7.3 | Inf | 5.696 | 12.93 | 0.000323 | 0.02616 | 1.58236 |
| MY18A_HUMAN | 12.3 | 0 | 12.3 | Inf | 6.39 | 12.4 | 0.0004287 | 0.0328 | 1.48413 |
